# Distinct nano-structures support a multifunctional role of actin at presynapses

**DOI:** 10.1101/2022.05.18.492480

**Authors:** Dominic Bingham, Florian Wernert, Julie Da Costa Moura, Fanny Boroni-Rueda, Nikki van Bommel, Ghislaine Caillol, Marie-Jeanne Papandréou, Christophe Leterrier

## Abstract

Synapses are the nexus of signal transmission in the nervous system. Despite decades of work, the architecture of the actin cytoskeleton that concentrates at presynapses remain poorly known, hindering our comprehensive understanding of its roles in presynaptic physiology. In this work, we take advantage of a validated model of bead-induced presynapses to measure and visualize isolated presynaptic actin by diffraction-limited and super-resolution microscopy. We first identify a major population of actin-enriched presynapses that concentrates more presynaptic components, and shows higher synaptic vesicle cycling than their non-enriched counterparts. Using pharmacological perturbations, we determine that an optimal amount of actin is necessary for this effect of actin enrichment. Modulation of this effect by actin nucleation inhibitors indicates its dependance on distinct presynaptic actin assemblies. Using Single Molecule Localization Microscopy (SMLM), we directly visualize these nano-structures in isolated presynapses, defining an actin mesh at the active zone, actin rails between the active zone and deeper reserve pools, and actin corrals around the whole presynaptic compartment. We finally show that these three types of presynaptic actin nano-structures are differentially affected by actin nucleation inhibitors, consistent with their effect on presynaptic component concentration and on synaptic vesicle cycling.

## Introduction

In the human brain and nervous system, hundreds of trillions (∼10^14^) of synapses ensure fast communication between neurons (Cano-Astorga et al., 2021; DeWeerdt, 2019). Chemical synapses are intricate assemblies connecting the emitting neuron, usually from a presynapse located along the axon, through a ∼20 nm synaptic cleft to the receiving neuron, typically along its dendrites. Following calcium elevation, synaptic vesicles fuse at the presynaptic active zone, releasing neurotransmitters through the cleft (Südhof, 2021). Since their discovery by pioneering electron microscopy (EM) works in the middle of the 20th century (Sotelo, 2020), the architecture of presynapses and their diversity have been progressively unraveled. Decade of studies have defined the presence and role of hundreds of specific synaptic proteins, culminating in elaborate molecular models of the presynaptic compartment (Wilhelm et al., 2014). Presynaptic boutons and terminals exhibit unique molecular specialization and structural plasticity, yet the detailed arrangement of the presynaptic cytoskeleton supporting these processes remains poorly known compared to their post-synaptic counterpart. This is particularly true for actin, despite its prominent enrichment in both pre- and postsynapses (Cingolani and Goda, 2008; Gentile et al., 2022).

The presence of actin at presynapses has been recognized from early EM studies, but its precise organization was difficult to pinpoint because the challenging size, lability and low contrast of actin filaments in EM preparations (Leterrier, 2021a; Papandréou and Leterrier, 2018). Short actin filaments are seen within the cytomatrix of the active zone, contacting closely apposed synaptic vesicles that form the readily-releasable pool (RRP, Hirokawa 1989, Li 2010). Actin filaments are also present deeper in the presynapse, where they are found around or within the vesicle clusters of the reserve pool, that can be mobilized by prolonged or intense stimulation (Landis and Reese, 1983; Siksou et al., 2007). In large presynapses of the lamprey, actin also forms structures between the periphery of the active zone, where endocytosis occurs for synaptic vesicle recycling, and the reserve pool (Bloom et al., 2003). The overall view of presynaptic actin organization from EM studies points to the existence of different nano-structures (Dillon and Goda, 2005), but is still too fragmented and diverse for a comprehensive architectural model to emerge. Fluorescence optical microscopy can provide a much better throughput and molecular specificity to visualize presynaptic components, but its limited spatial resolution (∼250 nm) precludes obtaining details within the ∼1 µm presynaptic bouton. Fortunately, super-resolution microscopy techniques can bypass this limitation, easily reaching resolutions down to tens of nanometers in fixed samples (Jacquemet et al., 2020). Single Molecule Localization Microscopy (SMLM) techniques, such as Stochastic Optical Reconstruction Microscopy (STORM) or DNA-Point Accumulation in Nanoscale Topography (PAINT) work by localizing single emitters in “blinking” samples, progressively building an image from single fluorophore coordinates with a resolution down to 10-15 nm (Lelek et al., 2021). In neurons, SMLM revealed a striking periodic organization of actin along the axon shaft: submembrane actin rings are regularly spaced every 190 nm by tetramers of spectrins (Leterrier, 2021b; Xu et al., 2013); this periodic actin-spectrin scaffold could only recently be visualized by electron microscopy (Vassilopoulos et al., 2019). At the presynapse, SMLM has been used to delineate the organization of specific components, but the organization of presynaptic actin has not been addressed so far (Carvalhais et al., 2021; Dani et al., 2010; Nosov et al., 2020). This is likely due to the challenge of densely labeling presynaptic actin (Reshetniak and Rizzoli, 2019) and distinguishing it from the high concentration and dense meshwork of postsynaptic actin that is evident in platinum-replica EM images of synapses (Korobova and Svitkina, 2010).

This lack of a comprehensive view on presynaptic actin architecture keeps us from understanding the roles of actin in presynaptic physiology and its functions in the various steps of the synaptic vesicle cycle: exocytosis, endocytosis, trafficking to and between vesicular pools. The diversity of synaptic types studied cannot alone explain the large span of different and often contradictory results on presynaptic functions obtained after perturbation of the actin cytoskeleton (Cingolani and Goda, 2008; Rust and Maritzen, 2015; Wu and Chan, 2022). Application of drugs that perturb actin assembly have led to reduced, unchanged or elevated vesicular release depending on the model, drug and experimental conditions used, leading to a number of conclusions on a driving, resisting or merely scaffolding role of presynaptic actin in the synaptic vesicle cycle (Bleckert et al., 2012; Cole et al., 2000; Li and Murthy, 2001; Morales et al., 2000; Rampérez et al., 2019; Sakaba and Neher, 2003; Sankaranarayanan et al., 2003). A plausible explanation of these contrasted results is that distinct actin structures co-exist within presynapses that have different functions in the synaptic vesicle cycle, with a net effect dependent on the precise experimental conditions (Cingolani and Goda, 2008; Papandréou and Leterrier, 2018). This further highlights the need to directly visualize and identify these presynaptic actin nano-structures, in order to generate meaningful hypotheses on their function.

In this work, we thus aimed at delineating and characterizing the presynaptic actin nano-structures. We first set up and validated a bead-induced model of presynapses in cultured hippocampal neurons that allowed us to distinguish presynaptic actin by both diffraction-limited and super-resolution microscopy. We could thus characterize the actin enrichment in a dominant subpopulation of induced presynapses and its effect on the presynapse composition and cycling activity. We assessed the effect of acute perturbations and modulations of the actin cytoskeleton on presynapses, validating induced presynapses against natural synapse in the same cultures. We then used super-resolution STORM and PAINT to directly visualize presynaptic actin nano-structures, demonstrating the existence of three main types of actin assemblies likely responsible for distinct functions within presynapses.

## Results

### Bead-induced presynapses allow to isolate presynaptic actin for imaging and quantification

The gold standard reagent to label filamentous actin in fixed neurons is fluorescent phalloidin (Melak et al., 2017). It provides a dense labeling of actin filaments, allowing for both quantification of the amount of actin on diffraction limited images, and visualization of the nanoscale actin structures in super-resolution microscopy (Jimenez et al., 2020; Lavoie-Cardinal et al., 2020). However, phalloidin stains all neurons in a fixed culture, labeling both the presynaptic and post-synaptic actin at synapses. As postsynapses are highly enriched in actin, notably within the dendritic spine heads (Allison et al., 1998), their close apposition hides the lower amount of presynaptic actin (Zhang and Benson, 2002) (Fig. S1). This is true on diffraction-limited images where pre- and post-synapse are not resolved (Fig. S1A, S1C), but also on STORM images, despite their ∼10X better resolution (Jacquemet et al., 2020). Furthermore, under STORM conditions optimized for the low amount of axonal actin, the highly enriched dendrites and spines are too densely labeled for single-molecule imaging, resulting in ill-defined structures that entirely mask the presynaptic actin structures (Fig S1B, S1D-E). Using a deep-learning algorithm with demonstrated performance in high-density labeling areas (Speiser et al., 2021), a clearer picture of both axonal and dendritic structures is obtained, but even this state-of-the-art approach does not allow to distinguish presynaptic from post-synaptic actin (Fig. S1F).

To visualize the specific enrichment and organization of presynaptic actin, we turned to a validated model of presynaptic induction using polylysine-coated beads (Lucido et al., 2009). As discovered more than 40 years ago (Burry, 1980), polylysine-coated beads seeded on neuronal cultures will induce the formation of presynaptic compartments within tens of minutes after contacting axons (Suarez et al., 2013). These induced specializations are bone fide presynapses: they show the structural hallmark of presynapses, including synaptic vesicle pools, are enriched in presynaptic protein components, and are competent for vesicular release (Burry, 1980; Burry et al., 1986; Lucido et al., 2009). They form by inducing recruitment of heparan sulfate proteoglycans at the contact site, including neurexins that bears an heparan sulfate glycan modification (Zhang et al., 2018). In recent years, this model of bead-induced presynapses has been used to dissect the mechanism of presynapse formation, in particular the need for local axonal translation (Batista et al., 2017; Taylor et al., 2013).

We thus seeded neurons after 7 to 8 days in vitro (div) with 5 µm-diameter polylysine-coated beads to induce isolated presynapses. After 48h, we fixed and immunolabeled neurons for presynaptic markers to check for presynaptic induction (Fig. 1). Fluorescence images show beads readily inducing a concentration of presynaptic components when contacting axons: scaffold proteins bassoon and synapsin (Fig. 1A-B) as well as synaptic vesicle proteins synaptophysin and vamp2 (Fig. 1C-D). 80 ± 2.3% of axon-contacting beads induce such presynaptic components concentration (S+ population, mean ± SEM, Fig. 1G), whereas 20 ± 3.3% of beads do not result in visible presynaptic specialization along the contacted axon (S– population) (see all statistics and tests for significance in Sup. File 1). We next assessed if the induced presynaptic specializations were competent for vesicular cycling using two methods. Firstly, an antibody against an extracellular epitope of synaptotagmin can accumulate in presynapses either after constitutive cycling during a prolonged incubation, or after stimulated cycling during a short incubation together with KCl (Kraszewski et al., 1995) (Fig. S2A). In our bead-seed-ed cultures, the anti-synaptotagmin antibody readily labels bead-induced presynaptic specializations (as detected by post-fixation synaptophysin labeling) in both the constitutive (Fig. 1E-F) and stimulated (Fig. S2B-C) versions of the assay. Secondly, we used the styryl dye FM1-43 to label the cycling synaptic vesicles in living neurons (Gaffield and Betz, 2006), loading then releasing the FM dye using two successive potassium chloride stimulations (Fig. S2D-E). FM dye readily accumulates in most enlarged contacts between axons and beads (as visualized using the live-cell actin probe SiR-actin) (Lukinavicius et al., 2014), and is released after the second stimulation (Fig. S2F).

**Figure 1:**
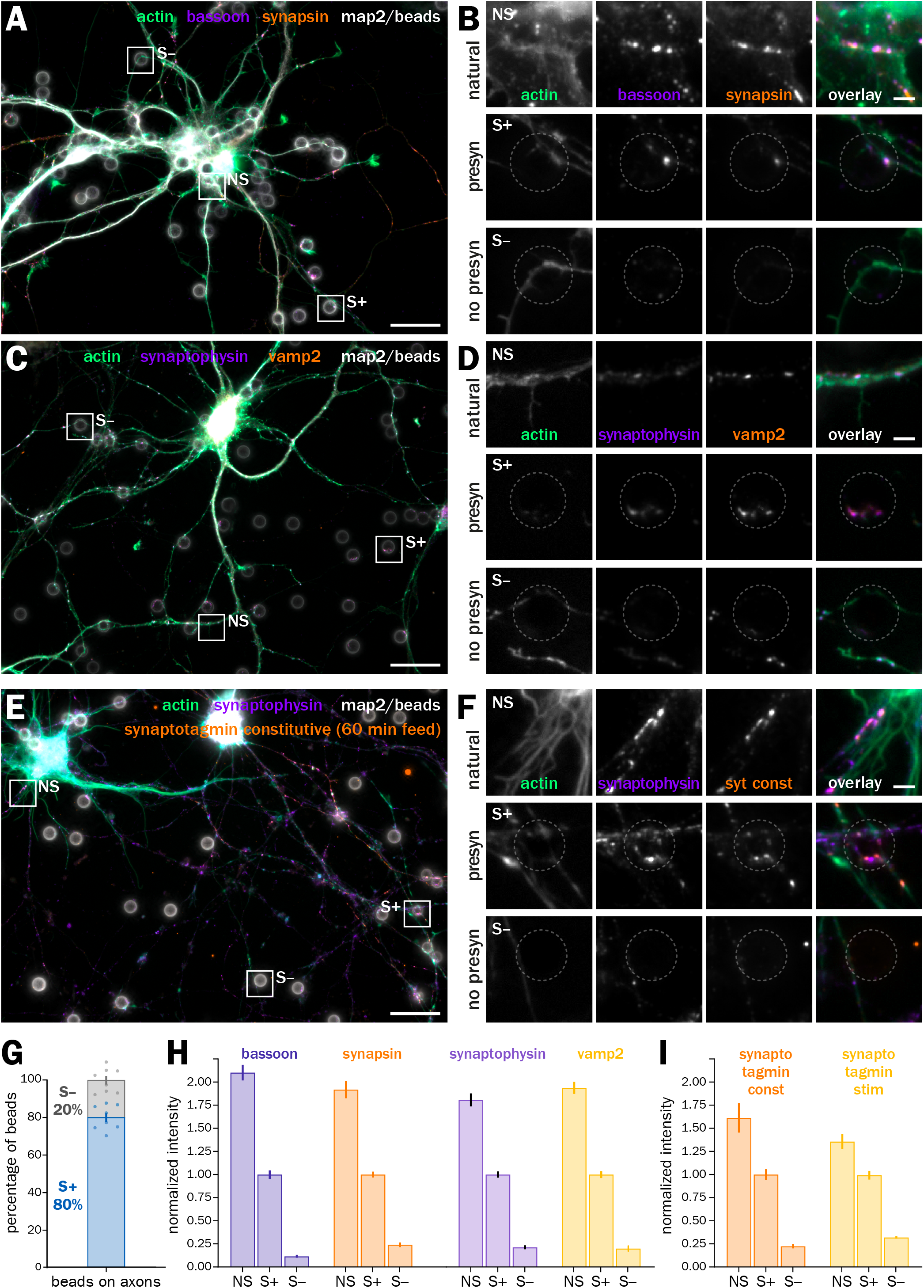
Polylysine-coated beads induce functional presynaptic specializations at axon-bead contacts. A. Widefield fluorescence image of cultured neurons seeded with polylysine-coated beads (gray) after 8 div and fixed 48h later at 10 div, labeled for actin (green), bassoon (purple), synapsin (orange), and map2 (gray). B. Zooms corresponding to the areas highlighted in A: top row, natural synapse at axon-dendrite contact (NS); middle row, induced presynapse at an axon-bead contact (S+); bottom row, axon-bead contact with no induced presynapse (S–). The position of the bead is indicated by the dashed gray circle. C. Widefield fluorescence image of cultured neurons 2 days after bead seeding at 8 div, labeled for actin (green), synaptophysin (purple), vamp2 (orange), and map2 (gray). D. Zooms corresponding to the natural synapse, S+ and S– axon-bead contacts highlighted in C. E. Widefield fluorescence image of cultured neurons 2 days after bead seeding at 8 div, labeled for actin (green), synaptophysin (purple), map2 (gray), and feeding with anti-synaptotagmin antibody (syt) during constitutive vesicular cycling (orange). F. Zooms corresponding to the natural synapse (NS), S+ and S– axon-bead contacts highlighted in E. Scale bars for A, C, E: 20 µm; for B, D, F: 2 µm. G. Quantification of the proportion of S+ (blue) and S– (gray) ax-on-bead contacts. H. Quantification of the labeling intensity for bassoon (purple), synapsin (orange), synaptophysin (light purple) and vamp2 (yellow) at natural synapses (NS), induced presynapses at axon-bead contacts (S+) and axon-bead contacts devoid of presynapse (S–), normalized to the intensity at S+ contacts. I. Quantification of the labeling intensity after syt feeding for constitutive (orange) and stimulated (KCL-induced, yellow) vesicular cycling at natural synapses (NS), induced presynapses at axon-bead contacts (S+) and axon-bead contacts devoid of presynapse (S–), normalized to the intensity at S+ contacts.

In these 9-10 div neurons 48h after beads seeding, we quantitatively assessed the enrichment of presynaptic components at axon-bead contacts where a presynaptic specialization has developed (S+), compared to the ones where it did not happen (S–), and to “natural” synapses between axons and dendrites in the same culture (NS, Fig. 1H). Intensity measurements of clusters at axon-bead contacts on fluorescence images show that induced presynaptic specializations S+ contain 4.8 - 10.6 times more bassoon, synapsin, synaptophysin and vamp2 than non-presynaptic contacts S–. This enrichment is not at high as in natural synapses, which have a 1.8 - 2.1 times higher cluster intensity than induced presynaptic specializations. This might be because a majority of natural synapses appear earlier than induced ones and accumulate more components, as synaptogenesis starts around 4-5 div in these cultures (Danielson et al., 2021; Fletcher et al., 1994; Green et al., 2019) whereas beads are present only for two days before fixation. In addition, the higher level of background and non-specific signal at dendrite-axon contacts might bias the identification of natural synapses toward brighter clusters. In addition, we measured the amount of vesicular cycling at axon-bead contacts using the anti-synaptotagmin (syt) antibody feeding assay: induced presynaptic specializations accumulate 2.6 and 4.0 times more syt antibody than non-presynaptic contacts, and 1.6 or 1.4 times less than natural synapses for constitutive or stimulated cycling, respectively (Fig. 1H). In conclusion, we confirmed that the polylysine bead-seeding model robustly induces isolated, functional presynapses in cultured hippocampal neurons, making it a valid model to discriminate presynaptic actin enrichment and organization.

### Actin is enriched in a major subpopulation of induced presynapses, which show elevated concentration in presynaptic components and higher synaptic cycling

Having validated our bead-induced presynapse model, we next focused on the actin content at induced presynapses (Fig. 2). Phalloidin labeling was visible along the axons and at all bead-induced presynapses. A majority of axon-bead contacts where a presynaptic specialization was induced (labeled S+ in Fig. 1) show specific enrichment in actin compared to the nearby axon shaft (Fig. 2A-D) and were labeled “actin +” (A+), whereas a minority of induced presynapses do not show actin enrichment and were labeled “actin –” (A–). A+ and A– induced presynapses are often found along the same axon arborization, suggesting it is not dependent on a cell-wide state (Fig. 2A-D). Quantitatively, 66 ± 3.4% of all S+ induced presynapses are A+ (actin-enriched), corresponding to 53 ± 2.0% of all axon-bead contacts (Fig. 2E). We measured the intensity of presynaptic components clusters at A+, A– and S– axon-bead contacts, to compare the concentration of components in A+ presynapses with A– presynapses and with non-presynaptic S– contacts (Fig. 2F). In all quantifications, we normalized the cluster intensities to the dominant A+ population. By definition, actin intensity is much lower in A– than A+ presynapses, similar to non-presynaptic contacts S– (0.25 ± 0.02 and 0.29 ± 0.03, respectively, Fig. 2F). The intensity of presynaptic component clusters in A– presynapses is significantly below that of A+ presynapses (below 1 after normalizing to A+ presynapses: 0.74 ± 0.03 for bassoon; 0.70 ± 0.03 for synapsin; 0.66 ± 0.03 for synaptophysin; 0.65 ± 0.03 for vamp2 Fig. 2F). Altogether, two thirds of induced presynapses are specifically enriched in actin, and this enrichment is associated with a 30 to 50% enrichment in all presynaptic components assessed.

**Figure 2:**
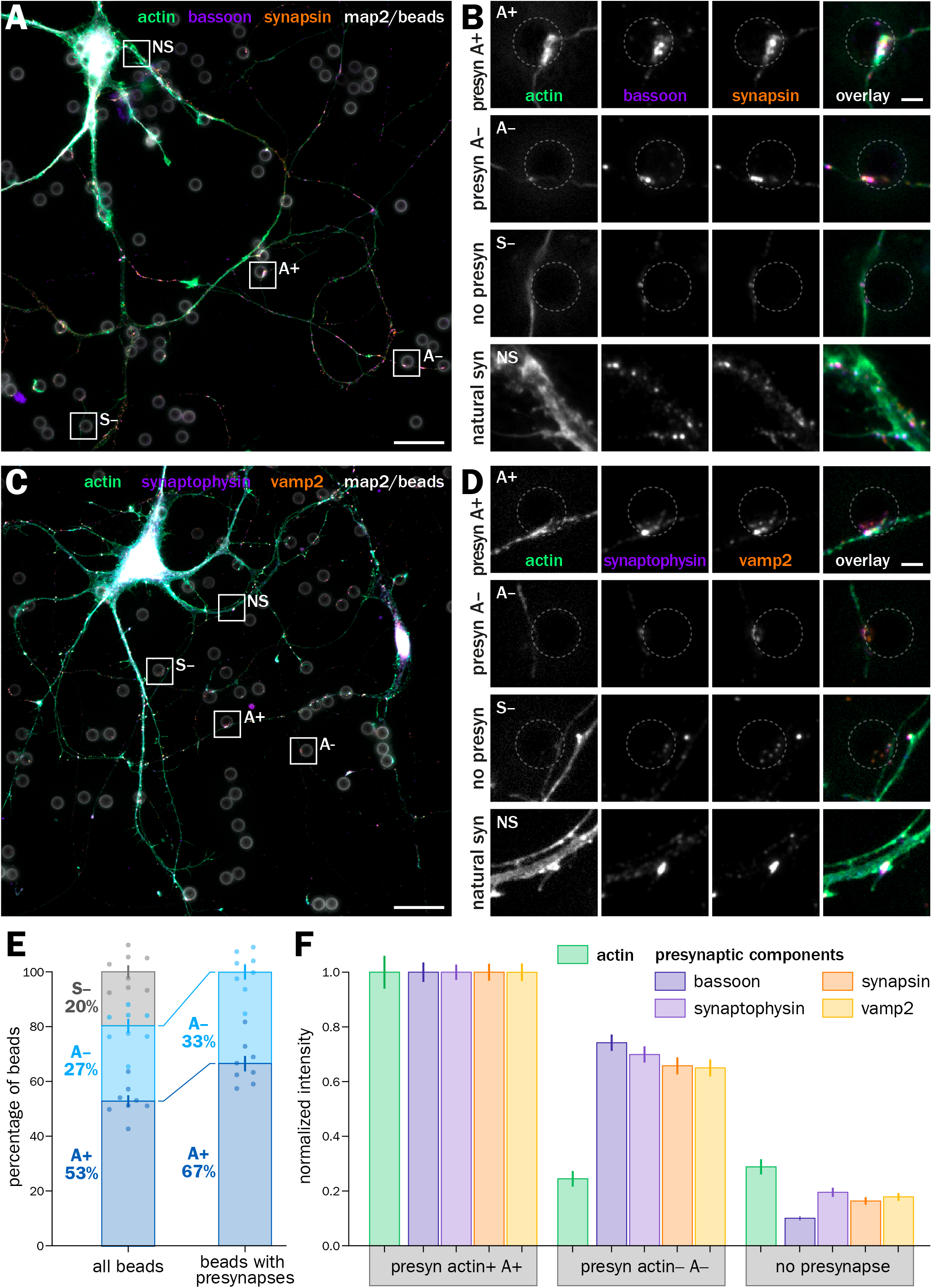
A major proportion of bead-induced presynapses are enriched in actin, and concentrate more presynaptic components than non-enriched induced presynapses. A. Widefield fluorescence image of cultured neurons 2 days after bead seeding at 8 div, labeled for actin (green), bassoon (purple), synapsin (orange), and map2 (gray). B. Zooms corresponding to the areas highlighted in A: top row, actin-enriched induced presynapse at an axon-bead contact (A+); second row, induced presynapse at an axon-bead contact with no actin enrichment (A–); third row, axon-bead contact with no induced presynapse (S–); bottom row, natural synapse at axon-dendrite contact (NS). C. Widefield fluorescence image of cultured neurons 2 days after bead seeding at 8 div, labeled for actin (green), synaptophysin (purple), vamp2 (orange), and map2 (gray). D. Zooms corresponding to the A+, A–, S– axon-bead contacts and natural synapses (NS) highlighted in C. Scale bars in A, C: 20 µm; B, D: 2 µm. E. Quantification of the proportion of A+ (dark blue), A– (light blue) and S– (gray) axon-bead contacts. F. Quantification of the labeling intensity for actin (green), bassoon (dark purple), synaptophysin (purple), synapsin (orange), and vamp2 (yellow) at actin-enriched presynapses (A+), induced presynapses with no actin enrichment (A–), and axon-bead contacts devoid of presynapse (S–), normalized to the intensity at A+ presynapses.

We then explored what are the factors for this selective enrichment of actin at induced presynapses. First, we tested if the maturity of seeded neurons could have an effect on actin enrichment at presynapses (Fig. S3). We seeded neurons during the third week in culture (polylysine beads seeded at 14 div and fixation at 16 div) instead of the second week. Classification of axon-bead contacts into A+, A– and S– and quantification of presynaptic clusters intensities show similar results to neurons seeded with beads between 8 and 10 days (Fig. S3A-D): 71 ± 2.5% of induced presynapses are enriched in actin (Fig. S3E), and these A+ presynapses have a significantly higher intensity of presynaptic components clusters (normalized intensities of Aclusters: 0.78 ± 0.05 for bassoon; 0.64 ± for synapsin; 0.76 ± 0.05 for synaptophysin; 0.83 ± 0.11 for vamp2, Fig. S3F). This shows that the maturation stage of neurons is not a key factor in the development of induced presynapses, and that second-week neurons are as competent as third-week ones for developing isolated presynapses.

Next, we tested if the overall proportion of A+ and A– induced presynapses is due to an underlying difference between presynapses induced along axons from excitatory (glutamatergic) and inhibitory (GABAergic) neurons. In cultures of embryonic rat hippocampal neurons, only 6% of neurons are GABAergic (Benson et al., 1994), but their synapses represent between 20 and 40% of all synapses (Danielson et al., 2021; Harms and Craig, 2005). We labeled bead-seeded neurons using markers for glutamatergic (VGLUT) and GABAergic (VGAT) presynapses (Fig. S4A-D) and measured the proportion of A+ vs A– induced presynapses as well as the intensity of presynaptic component clusters in each population. Actin-enriched presynapses are similarly dominant in the two synapse types (66 ± 8% and 79 ± 6% of A+ presynapses for glutamatergic and GABAergic presynapses, respectively). In both synapse types, actin enrichment is associated with an elevated concentration of presynaptic components (including VGLUT and VGAT), with normalized cluster intensities below 1 for the A– presynapses (glutamatergic presynapses: 0.49 ± 0.09 for synapsin and 0.57 ± 0.08 for VGLUT; GABAergic presynapses: 0.42 ± 0.07 for synapsin and 0.70 ± 0.12 for VGAT). This shows that the two populations of actin-enriched and non-enriched presynapses are similarly present in excitatory and inhibitory presynapses, suggesting a general phenomenon that does not depend on the presynapse type.

Finally, we assessed whether actin enrichment in an induced presynapse is associated with a difference in synaptic vesicle cycling. It has been shown that actin was involved in activating silent synapses after stimulation, suggesting that non-en-\riched presynapses could even be silent in our cultures (Yao et al., 2006). We used the syt antibody feeding assay (Fig. S2A) to measure the constitutive and stimulated vesicular cycling at actin-enriched and non-enriched induced presynapses, while monitoring their synaptophysin concentration in the same experiment (Fig. 3A-D). A+ presynapses show a 20 to 40% more intense syt accumulation than A– presynapses, with the A– normalized intensity of syt clusters being below 1 for both constitutive and stimulated version of the assay (0.82 ± 0.08 for constitutive cycling, non-significant difference with A+; 0.72 ± 0.03 for stimulated cycling, Fig. 3E). We confirmed these results using FM1-43 staining experiments in living neurons co-stained with SiR-actin to reveal the actin enrichment at induced presynapses, measuring FM1-43 intensity after dye loading and release by two successive short KCl incubations (Fig. 3F). Actin-enriched presynapses A+ could accumulate more FM1-43 than non-actin enriched ones A– that had a lower normalized intensity (0.71 ± 0.07), while the intensity level after dye release was similarly low (0.38 ± 0.03 for A+ and 0.32 ± 0.04 for A– after release, normalized to the loaded A+ intensity, Fig. 3G). Thus, lack of actin enrichment is not associated with silent presynapses, but actin-enriched presynapses tend to have a significantly more robust vesicular cycling activity.

**Figure 3:**
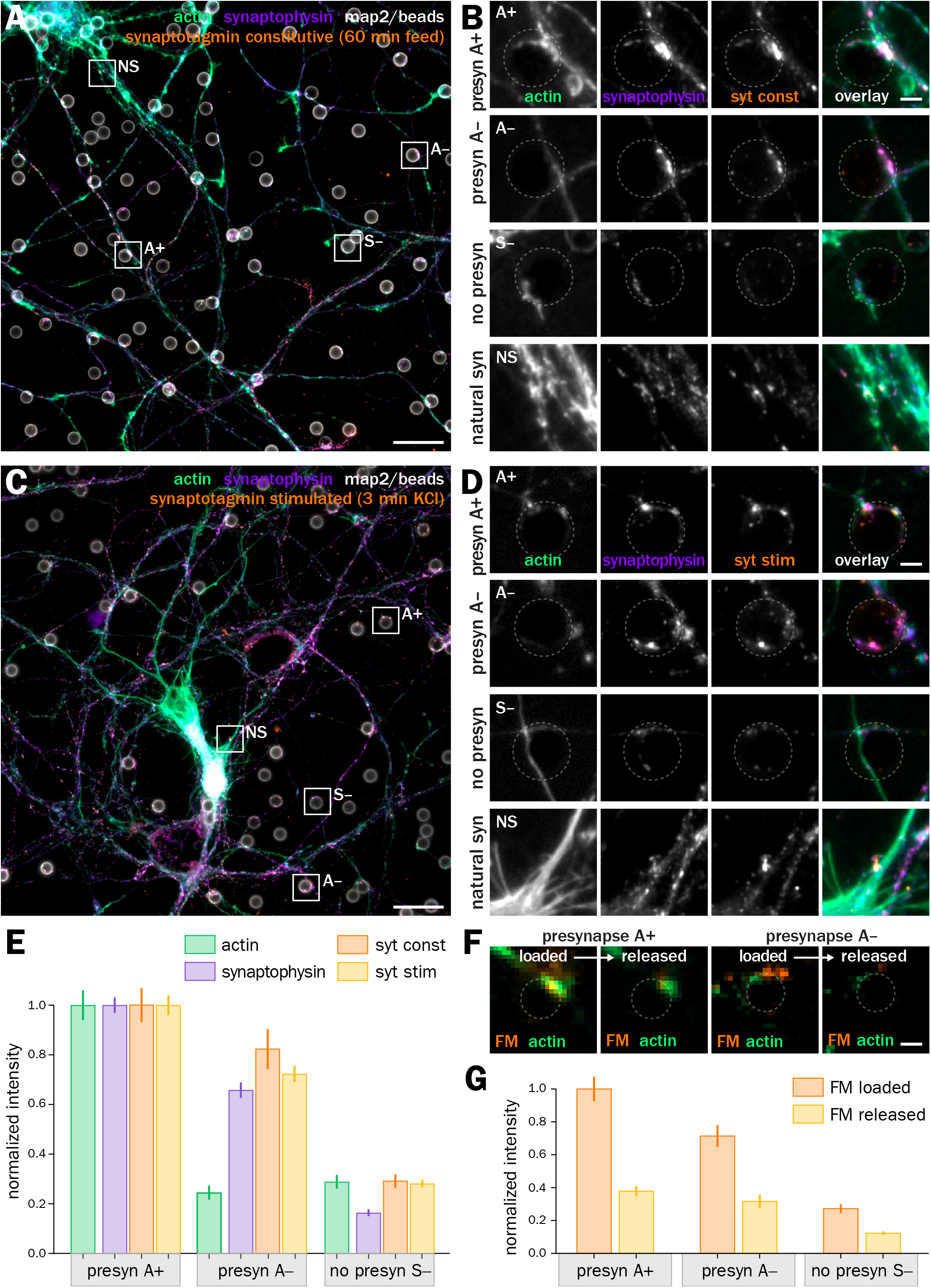
Actin-enriched induced presynapses have a higher vesicular cycling than non-enriched induced presynapses. A. Widefield fluorescence image of cultured neurons 2 days after bead seeding at 8 div, labeled for actin (green), synaptophysin (purple), synapsin (orange) and map2 (gray). B. Zooms corresponding to the A+, A–, S– axon-bead contacts and natural synapses (NS) highlighted in B.C. Widefield fluorescence image of cultured neurons 2 days after bead seeding at 8 div, labeled for actin (green), synaptophysin (purple), and map2 (gray) and feeding with anti-synaptotagmin antibody (syt) during stimulated vesicular cycling. D. Zooms corresponding to the A+, A–, S– axon-bead contacts and natural synapses (NS) highlighted in C. Scale bars in A, C, 20 µm; B, D, 2 µm. E. Quantification of the labeling intensity for actin (green), synaptophysin (purple), and after syt feeding for constitutive (orange) and stimulated (yellow) vesicular cycling at actin-enriched presynapses (A+), induced presynapses with no actin enrichment (A–), and axon-bead contacts devoid of presynapse (S–), normalized to the intensity at A+ presynapses. F. Zooms on axon-bead contacts of living neurons stained with SiR-actin (green) and loaded with FM1-43 (orange, left image) before release (right image). Left two images show an A+ presynapse, right ones an A– presynapse. Scale bar, 2 µm. G. Quantification of the FM staining intensity after loading (orange) and release (yellow) in A+ and A– induced presynapses as well as axon-bead contacts devoid of a presynapse (S–).

### Both acute disassembly and over-stabilization of actin content lower the concentration of presynaptic components and vesicular cycling in presynapses

To test if actin enrichment was responsible for the higher concentration of presynaptic components and higher cycling rate of A+ presynapses, we perturbed actin using drugs able to either disassemble and sever actin filaments (swinholide A) (Spector et al., 1999) or inhibit their depolymerization and over-stabilize them (cucurbitacin E) (Sörensen et al., 2012). We used short, acute perturbations with strong drugs in order to avoid indirect effects from impacting other aspects of the neuronal physiology (Fig. 4) (Vassilopoulos et al., 2019). 1h treatment with 1 µM swinholide A (swin) results in the near-complete disappearance of filamentous actin at induced presynapses, as evidenced comparing phalloidin staining between the control and swin conditions (Fig. 4A-B), with all treated presynapses being classified as A– and the normalized intensity dropping to 0.02 ± 0.01 (Fig. 4D). Conversely, phalloidin staining is strongly elevated after treatment with 5 µM cucurbitacin E (cuc, Fig. 4C), which caused all treated presynapses to be classified as A+, and a rise of the normalized intensity to 7.3 ± 0.43 (Fig. 4E).

**Figure 4:**
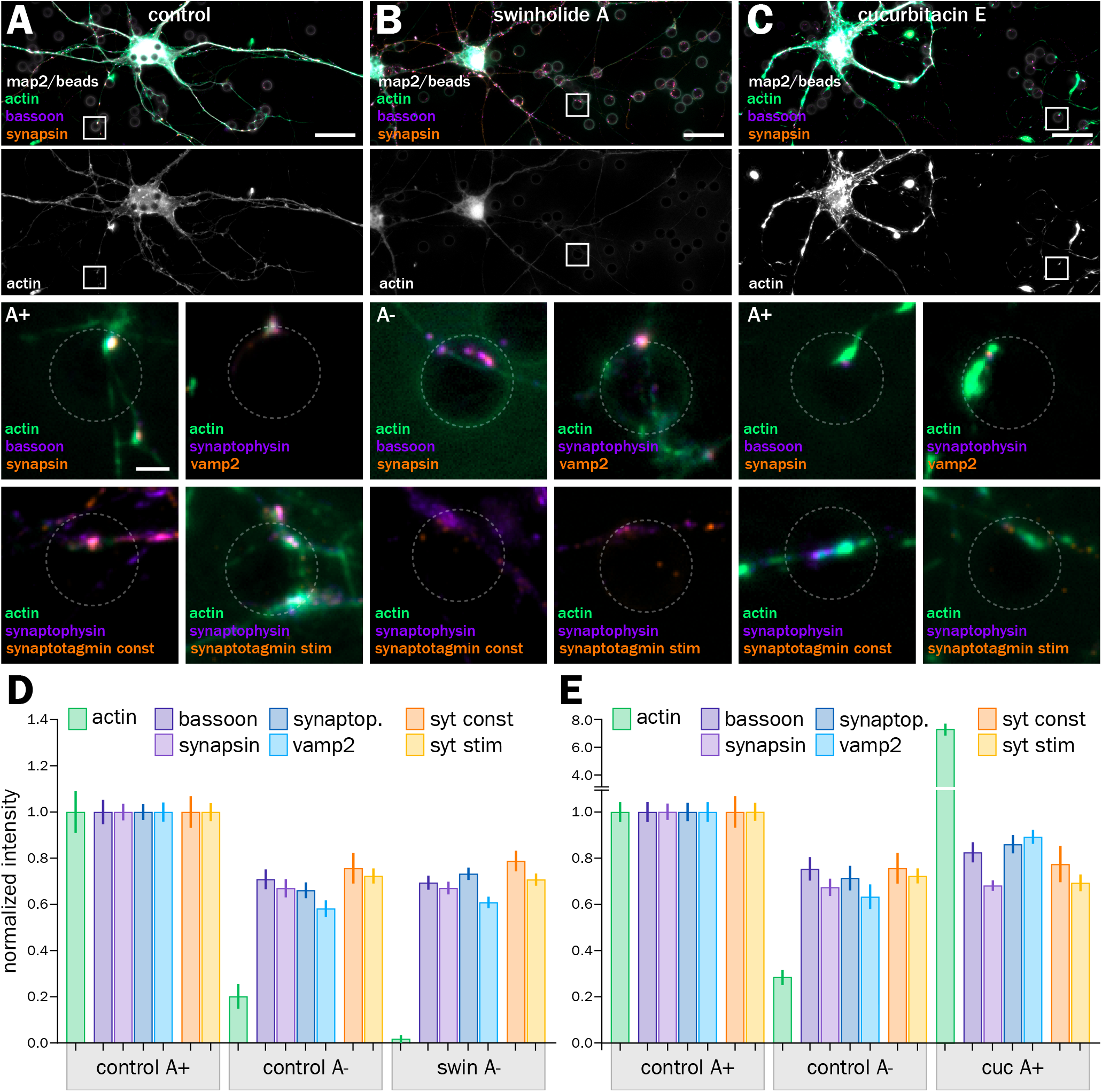
Acute actin disassembly and over-stabilization lowers the concentration of presynaptic components at induced presynapses. A. Top, widefield fluorescence image of cultured neurons 2 days after bead seeding at 8 div, control treated with vehicle (DMSO) for 1 h and labeled for actin (green), bassoon (purple), and map2 (gray). Bottom, panels showing A+ presynapses labeled for actin, presynaptic components and after syt feeding. The first panel is a zoom of the area highlighted in the top image. B. Top, widefield fluorescence image of cultured neurons 2 days after bead seeding at 8 div, treated with 1 µM swinholide A (swin) for 1 h, and labeled for actin (green), bassoon (purple), and map2 (gray). Bottom, panels showing A– presynapses labeled for actin, presynaptic components and after syt feeding. The first panel is a zoom of the area highlighted in the top image. C. Top, widefield fluorescence image of cultured neurons 2 days after bead seeding at 8 div, treated with 5 µM cucurbitacin E (cuc) for 3 h, and labeled for actin (green), bassoon(purple), and map2 (gray). Bottom, panels showing A+ presynapses labeled for actin, presynaptic components and after syt feeding. The first panel is a zoom of the area highlighted in the top image. Scale bars on large images in A-C, 20 µm; on zoomed images, 2 µm. D. Quantification of the labeling intensity for actin (green), bassoon (dark purple), synapsin (purple), synaptophysin (dark blue), vamp2 (blue), and after syt feeding for constitutive (orange) and stimulated (yellow) vesicular cycling at actin-enriched presynapses (A+) and induced presynapses with no actin enrichment (A–) in the control condition, and at A– presynapses after swin treatment. E. Quantification of the labeling intensity for actin (green), bassoon (dark purple), synapsin (purple), synaptophysin (dark blue), vamp2 (blue), and after syt feeding for constitutive (orange) and stimulated (yellow) vesicular cycling at actin-enriched presynapses (A+) and induced presynapses with no actin enrichment (A–) in the control condition, and at A+ presynapses after cuc treatment.

We then measured the concentration of presynaptic components and vesicular cycling after swin and cuc treatments. Swin treatment decreased the concentration of all labeled components (normalized intensities of 0.69 ± 0.03 for bassoon, 0.67 ± 0.03 for synapsin, 0.74 ± 0.03 for synaptophysin, and 0.61 ± 0.03 for vamp2, Fig. 4D) to values very similar to those from untreated A– presynapses (Fig. 4D). In addition, actin disassembly by swin also reduced the vesicular cycling in both constitutive and KCl-stimulated assays (syt normalized intensities of 0.79 ± 0.04 for constitutive, 0.71 ± 0.03 for stimulated) to values similar to control A– presynapses (Fig. 4D). Interestingly, elevated polymerization and stabilization of actin by cuc also reduces presynaptic components concentration and vesicular cycling at induced presynapses (normalized intensities of 0.83 ± 0.04 for bassoon, 0, 68 ± 0.02 for synapsin, 0.86 ± 0.04 for synaptophysin, 0.89 ± 0.03 for vamp2, 0.77 ± for syt constitutive, 0.69 ± 0.04 for syt stimulated), to values similar to untreated A– presynapses (Fig. 4E).

This drop after swin treatment implies that in control presynapses, actin enrichment is necessary for the extra concentration of 30-50% in presynaptic components and vesicular cycling. Moreover, over-stabilization for actin also results in a drop of presynaptic components concentration and vesicular cycling, suggesting that an optimal level of presynaptic actin enrichment is needed to ensure their efficient functioning. To further confirm the validity of these findings, we measured the concentration of presynaptic components and the amount of vesicular cycling in natural synapses (dendrite-axon contact) on the same images as the bead-induced presynapses. In this case, the actin intensity measurement reflects the total actin at the synapse, and is likely dominated by postsynaptic actin. Actin disassembly by swin and over-stabilization by cuc leads to a 10-40% drop in presynaptic component concentration and vesicular cycling, perfectly in line with the effects observed in bead-induced presynapses (Fig. S5A-B).

### Inhibition of actin nucleators results in different effects on actin and presynaptic components in induced presynapses, suggesting action on distinct presynaptic actin structures

The fact that both actin disassembly and over-stabilization lead to a less efficient vesicular cycling is consistent with the idea that distinct actin structures coexist within a presynapse, and can have opposing effects on presynaptic function. To address this hypothesis more finely, we used drugs that target distinct actin filament nucleation mechanisms: CK666 inhibits Arp2/3-mediated branched actin nucleation (Nolen et al., 2009), whereas SMIFH2 inhibits the nucleation of linear actin structures mediated by formins (Rizvi et al., 2009). We thus evaluated the effect of CK666 and SMIFH2 treatments on actin-enriched induced presynapses (A+, Fig. 5A-C). 1h treatment with 50 µM CK666 results in a contrasted modulation of presynaptic components (Fig. 5D): total filamentous actin in A+ presynapses is not significatively affected (normalized intensity of 0.95 ± 0.04), indicating a shift of actin polymerization toward other nucleation mechanisms. Scaffold proteins bassoon and synapsin are downregulated (0.89 ± 0.04 for bassoon, 0.77 ± 0.04 for synapsin), whereas synaptic vesicle proteins are not significantly affected (1.07 ± 0.04 for synaptophysin and 1.04 ± 0.05 for vamp2). Interestingly, a tendency for higher constitutive and stimulated vesicular cycling is found, although non-significant (1.05 ± 0.09 for syt constitutive, 1.08 ± 0.07 for syt stimulated). Treatment with 30 µM SMIFH2 for 1h results in a significantly higher amount of actin in A+ presynapses (normalized intensity1.34 ± 0.06), but this compensation is associated with a general downregulation of presynaptic components (0.86 ± 0.04 for bassoon, 0.80 ± 0.05 for synapsin, 0. 71 ± 0.03 for synaptophysin, 0.89 ± 0.04 for vamp2) and a significant drop of vesicular cycling (0.55 ± 0.03 for syt constitutive, 0.73 ± 0.04 for syt stimulated, Fig. 5D).

**Figure 5:**
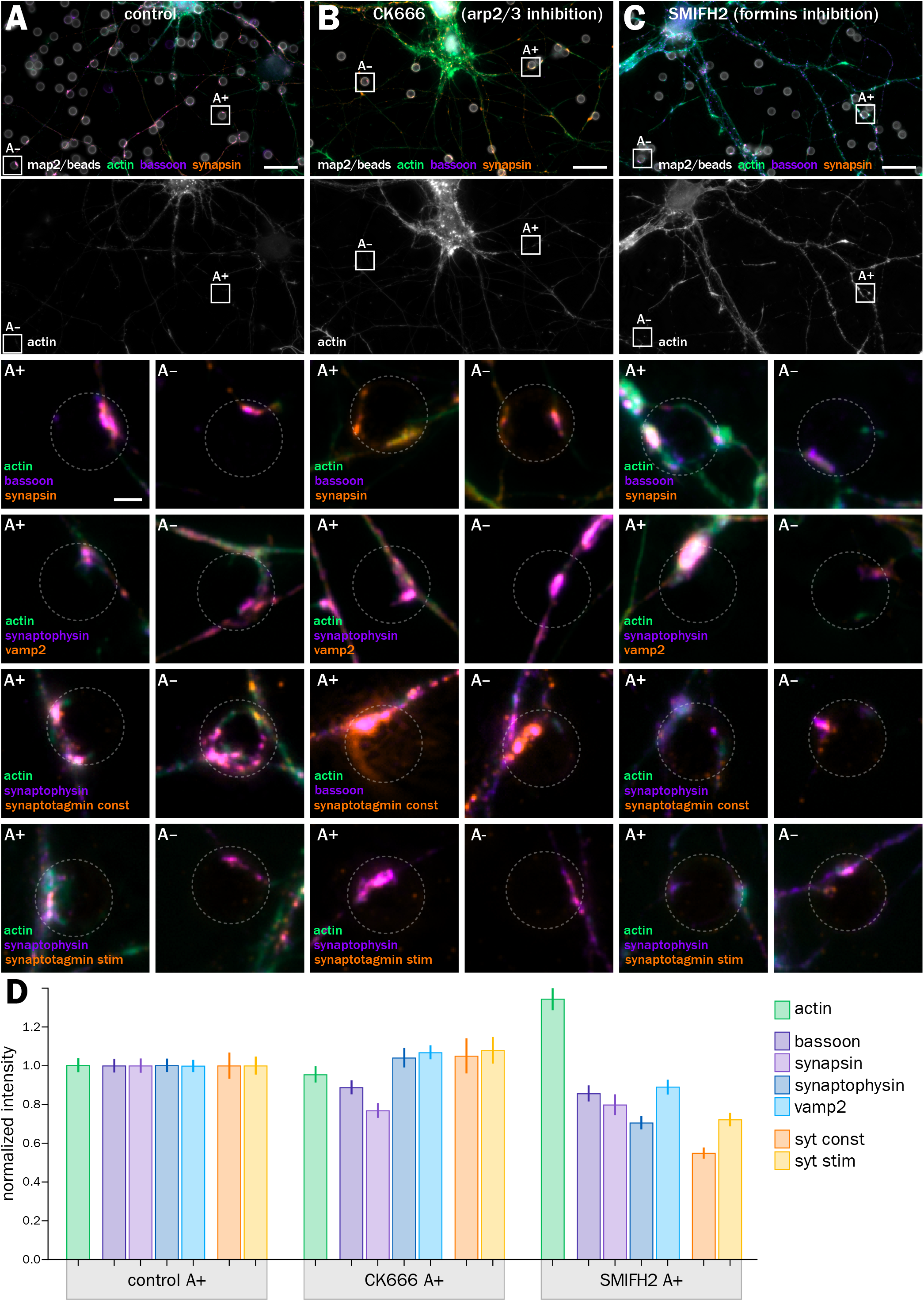
Arp2/3 and formins inhibitors have distinct effects on presynaptic components at induced presynapses. A. Top, widefield fluorescence image of cultured neurons 2 days after bead seeding at 8 div, control treated with vehicle (DMSO) for 1 h and labeled for actin (green), bassoon (purple), and map2 (gray). Bottom, panels showing A+ and A– presynapses labeled for actin, presynaptic components and after syt feeding. The top panels are zooms of the areas highlighted in the top image. B. Top, widefield fluorescence image of cultured neurons 2 days after bead seeding at 8 div, treated with 50 µM CK666 (Arp2/3 inhibitor) for 1 h, and labeled for actin (green), bassoon (purple), and map2 (gray). Bottom, panels showing A+ and A– presynapses labeled for actin, presynaptic components and after syt feeding. The top panels are zooms of the areas highlighted in the top image. C. Top, widefield fluorescence image of cultured neurons 2 days after bead seeding at 8 div, treated with 30 µM SMIFH2 (formins inhibitor) for 3 h, and labeled for actin (green), bassoon(purple), and map2 (gray). Bottom, panels showing A+ and A– presynapses labeled for actin, presynaptic components and after syt feeding. The top panels are zooms of the areas highlighted in the top image. Scale bars on large images in A-C, 20 µm; on zoomed images, 2 µm. D. Quantification of the labeling intensity for actin (green), bassoon (dark purple), synapsin (purple), synaptophysin (dark blue), vamp2 (blue), and after syt feeding for constitutive (orange) and stimulated (yellow) vesicular cycling at actin-enriched presynapses (A+) in the control condition, and after CK66 or SMIFH2 treatment.

By contrast, CK666 has no effect on induced presynapses non enriched in actin, and SMIFH2 has a limited effect on synapsin and synaptophysin content, as well as KCl-stimulated cycling (Fig. S5C). Here again, we verified the effects found on induced presynapses by quantifying the effect of CK666 and SMIFH2 on natural synapses in the same neuronal culture (Fig. S5D). The same treatments result in small variations that are similar but not significantly different from control for presynaptic components concentration, suggesting a tighter control or higher stability of the presynaptic compartment in natural synapses. Of note, vesicular cycling is strongly affected by inhibiting formins with SMIFH2, similarly to induced presynapses (Fig. S5D). Overall, the effect of CK666 and SMIFH2 on induced presynapses content and function reveal some insight on the role of different actin structures: small, contrasted effects of inhibiting Arp2/3 likely result from the inhibition of several different branched, Arp2/3-dependant actin structures, whereas the global downregulation observed with SMIFH2 suggests the presence of linear, formin-dependent actin structures that favor synaptic cycling.

### STORM of actin at induced presynapses reveal nano-structures with distinct sensitivity to actin nucleators inhibition

At this point, the presynaptic content in presynapses and its regulation by actin perturbations are consistent with the existence of distinct actin structures with different roles within presynapses. However, these inferred actin structures remain undiscernible due to the small size of the presynapses and the limited resolution of fluorescence microscopy. Taking advantage of bead-induced presynapses that isolate presynaptic actin, we used STORM to directly visualize actin nano-structures within presynapses with a lateral resolution of around 15 nm (Fig. 6) (Leterrier et al., 2015). In bead-seeded neuronal cultures, STORM exquisitely resolves phalloidin-labeled actin in both natural axon-dendrite synapses and in bead-induced presynapses enriched in actin (Fig. 6 A-B). As discussed above (see Fig. S1), the close apposition and high concentration of postsynaptic actin make discerning presynaptic actin in natural synapses impossible (Fig. 6B, top). By contrast, STORM clearly resolve presynaptic actin at axon-bead presynapses (Fig. 6B, bottom and 6C). The 190-nm spaced actin rings found along the axon shaft (Leterrier, 2021b) usually stop at the presynaptic bouton, confirming previous super-resolved spectrin images suggesting that boutons are devoid of the axonal actin/spectrin periodic scaffold (He et al., 2016; Sidenstein et al., 2016). However, the periodic scaffold sometimes persist along the membrane dorsal to the presynaptic contact, consistent with the idea that en-passant boutons are laterally apposed to the continuous axonal shaft. In addition, the vicinity of presynapses often shows fine longitudinal actin bundles, reminiscent of the actin trails that can transport actin and presynaptic components between adjacent boutons (Fig. 6C) (Chenouard et al., 2020; Ganguly et al., 2015).

**Figure 6:**
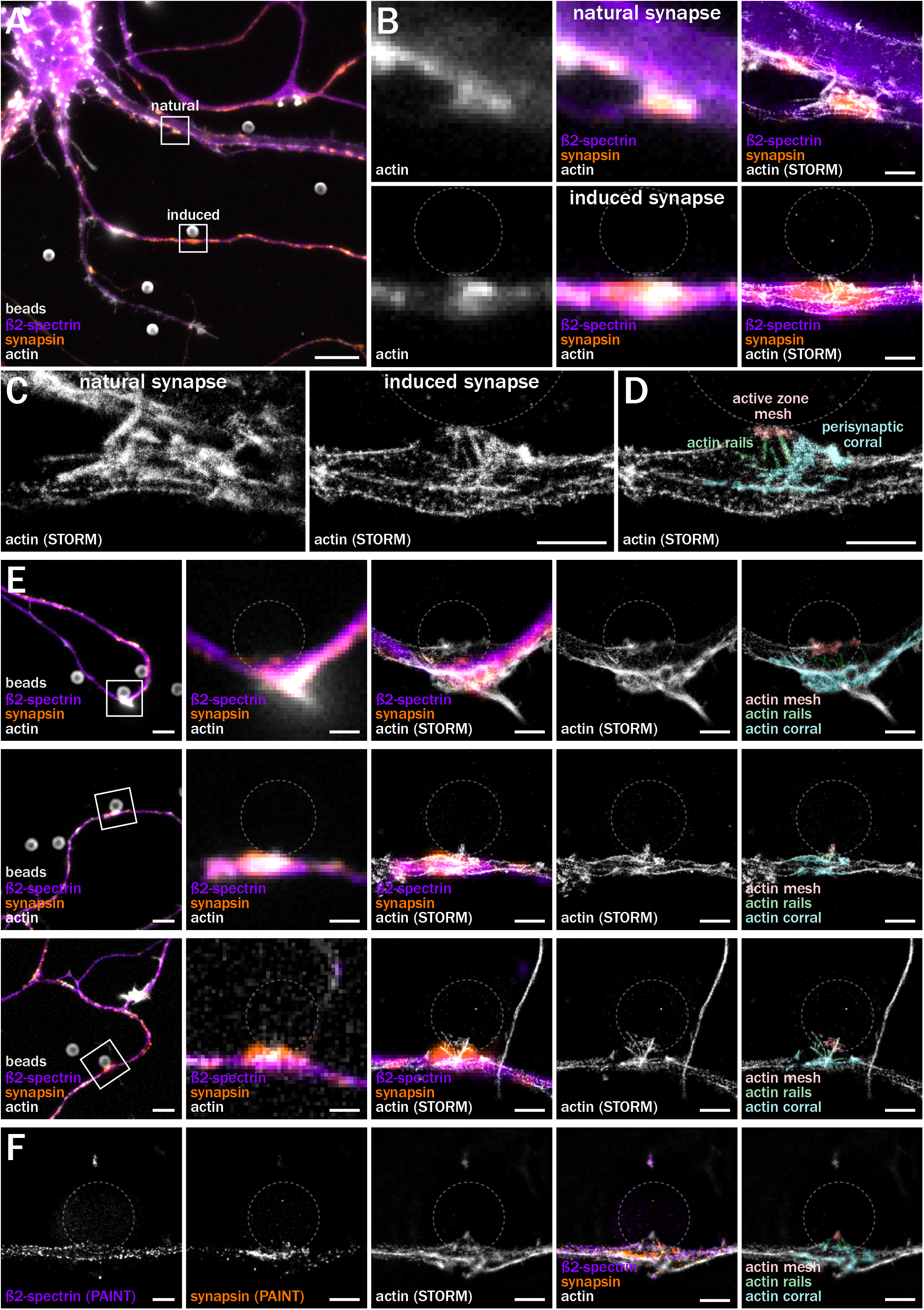
STORM visualizes distinct actin nano-structures at induced presynapses. A. Widefield fluorescence image of cultured neurons 2 days after bead seeding at 8 div, labeled for ß2-spectrin (purple), synapsin (orange) and actin (gray). B. Zooms on a natural synapse at an axon-dendrite contact (top row) and on an induced presynapse at an axon-bead contact (bottom row), corresponding to the highlighted areas in A. Left column are widefield images of actin (gray), middle channel are overlays of widefield images of ß2-spectrin (purple), synapsin (orange), and actin (gray); right column are overlays of widefield images of ß2-spectrin (purple) and synapsin (orange) with the STORM image of actin (gray). C. Zoomed STORM images of the natural synapse (left) and bead-induced presynapse (right). D. Same image as C, left, with color highlighting of presynaptic actin structures: active zone mesh (red), actin rails (green), perisynaptic corral (blue). E. Additional representative images of bead-induced presynapses. First column shows the widefield image of ß2-spectrin (purple), synapsin (orange) and actin (gray). Second column is a zoom of the highlighted area in the first image. Third column shows an overlay of widefield images of ß2-spectrin (purple) and synapsin (orange) with the STORM image of actin (gray). Fourth column is the isolated STORM image of actin (gray). Fifth column is the STORM image of actin with highlighted presynaptic actin structures: active zone mesh (red), actin rails (green), perisynaptic corral (blue). F. 3-color SMLM image of a bead-induced presynapse labeled for ß2-spectrin (PAINT image), synapsin (PAINT image) and actin (STORM image). The third column is an overlay of the three channels, while the fourth column shows the actin STORM image with highlighted presynaptic actin structures. Scale bars in A, 10 µm; in B-D, 1 µm; in C, 5 µm (first column) and 1 µm (other columns); in F, 1 µm.

Within induced presynapses themselves, STORM allowed us to delineate distinct nano-structures that we could classify in three types (Fig. 6D): a fine actin mesh at the bead contact, corresponding to the localization of the active zone (in red on Fig. 6D); fine, linear actin rails often connecting the active zone to deeper compartments within the bouton (green on Fig. 6D); and larger, branched structures surrounding the whole presynapse (synapsin-positive area) that we called actin corrals (in cyan on Fig. 6D). Similar to natural presynapses, induced presynapses show a significant variability in size, shape and architecture. Despite this, the three types of presynaptic actin structures (mesh, rails and corrals) were consistently identified in most of the 66 presynapses imaged by STORM, with the same relative arrangement to the point of bead contact (Fig. 6E). To refine the relationship between actin and presynaptic components, we performed 3-color STORM/PAINT acquisition, imaging actin with STORM before performing 2-color DNA-PAINT (Jimenez et al., 2020) for ß2-spectrin, which forms rings along the axon shaft, and synapsin, which delineates the pools of synaptic vesicles. This confirmed that the spectrin submembrane scaffold stops at the bouton, and that the actin corral encases the synaptic vesicle cluster, with rails pointing to the active zone (Fig. 6F).

STORM thus confirmed that, as inferred from pharmacological experiments, different actin structures coexist within presynapses. The nanoscale texture of the mesh and corrals suggest that they are made of branched actin filaments, while the rails are made of linear filaments. This means that they could be differentially regulated by Arp2/3 and formins inhibitors, supporting the effects we found for these inhibitors on presynaptic components concentration and vesicular cycling (see Fig. 5). To test this, we performed 3D-STORM of induced presynapses after a short treatment of bead-seeded cultures with CK666 (50 µM, 1h) and SMIFH2 (30 µM, 1h). We used deep learning-based DECODE processing (Speiser et al., 2021) to obtain a higher number of more precise localizations, resulting in better defined structures in 3D that we could identify as actin mesh, rails and corrals, blindly scoring each presynapse in a semi-quantitative manner (Fig. 7A-C). Inhibition of Arp2/3 mediated branched actin nucleation with CK666 resulted in a lower proportion of presynapses containing actin mesh (from 71% to 62% of presynapses) and actin corrals (from 69% to 62%), whereas it led to slightly more presynapses with actin rails (from 46% to 54%, Fig. 7D). Inhibition of formins-mediated linear actin nucleation minimally elevated meshes and corrals (71% to 78% and 69% to 72%, respectively), but led to a larger drop in the proportion of presynapses containing actin rails (46% to 33%). Although these effects of short term, acute treatments are partial and the semi-quantitative evaluation precludes assessing their significance, they are consistent with a distinct nature of the newly identified presynaptic actin structures.

**Figure 7:**
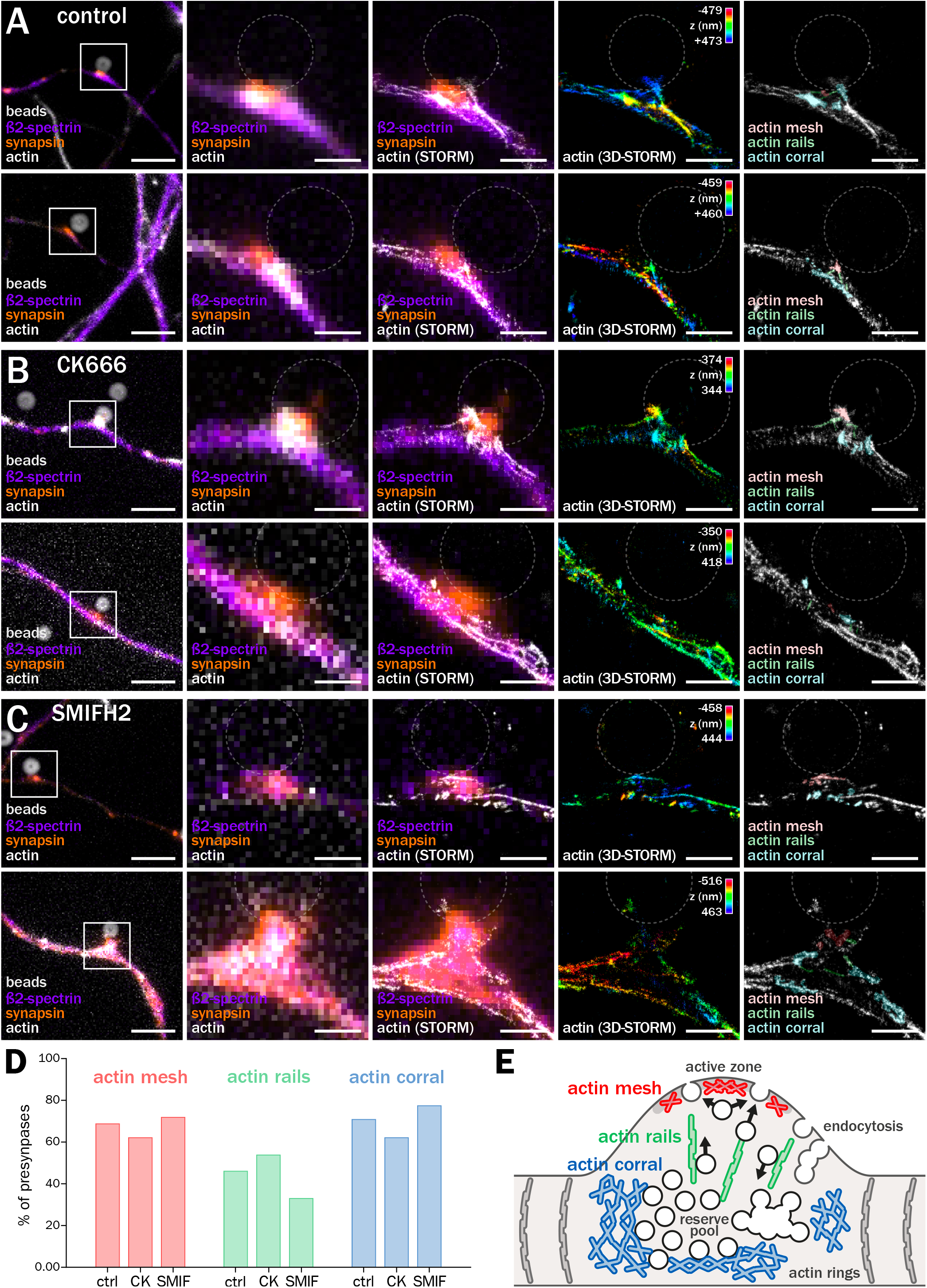
Presynaptic actin nano-structures are distinctly sensitive to Arp2/3 and formins inhibition. A-C. Images of induced presynapses from cultured neurons 2 days after bead seeding at 8 div. First column shows a widefield image of ß2-spectrin (purple), synapsin (orange) and actin (gray). Second column is a zoom of the highlighted area in the first column image. Third column shows an overlay of widefield images of ß2-spectrin (purple) and synapsin (orange) with the STORM image of actin (gray). Fourth column is the isolated 3D-STORM image of actin, color-coded for depth (with Z scale in the top right corner). Fifth column is the STORM image of actin with highlighted presynaptic actin structures: active zone mesh (red), actin rails (green), perisynaptic corral (blue). A, two representative examples from neuronal cultures treated with DMSO for 1 h (control condition). B, two examples after treatment with 50 µM CK666 for 1 h. C, two examples after treatment with 30 µM SMIFH3 for 1 h. Scale bars, 5 µm (first column) and 1 µm (other columns). D. Semi-quantitative assessment of the frequency of identification for presynaptic actin structures: actin mesh, rails and corrals, from all the presynapses imaged in the control condition (ctrl) or after treatment with CK666 (CK) or SMIFH2 (SMIF). E. Cartoon summarizing the architecture and role of the identified presynaptic actin structures: the branched actin mesh at the active zone (red) regulates access to the plasma membrane for exocytosis; linear actin rails (green) help vesicles move between the exocytosis, endocytosis zones and intracellular pools; the perisynaptic actin corrals (blue) scaffold the vesicular pool and could also help at endocytic zones.

If we bring together the first direct visualization of distinct Arp2/3 and formins-dependent presynaptic actin nano-structures (Fig. 6-7) with the effect of their modulation on synaptic components concentration and vesicular cycling (Fig. 5), we obtain a new view for the role of actin at presynapses (Fig. 7E). Three main actin structures are present within presynapses: an active zone mesh, rails between the active zone/ plasma membrane and the deeper reserve pools, and an actin corral encasing the reserve pool. Perturbation of branched actin structures via Arp2/3 inhibition destabilizes both the mesh and the corral, and the contrasted consequences (Fig.5D and S5C-D) indicate that these two structures have opposing roles in organizing vesicular cycling: for example, the submembrane mesh could be a resistive structure for exocytosis at the active zone, while the corrals have a scaffolding role positively contributing to reserve pool maintenance or endocytosis from around the active zone. Perturbation of the linear actin rails via formins inhibition has a net negative role on vesicular cycling, consistent with their possible role in helping the transport of synaptic vesicles between the reserve pool and active zone, or the shuttling of retrieved vesicles from their endocytosis point to the intra-synaptic pools (Fig 7E).

## Discussion

In this work, we addressed a long-standing gap in our understanding of the neuronal architecture by delineating and characterizing the presence of actin nano-structures within presynapses. We first made presynaptic actin visible by inducing isolated presynapses on polylysine-coated beads. This revealed that around two thirds of presynapses are enriched in actin, this enrichment being associated with a higher concentration of presynaptic components (scaffolds and synaptic vesicle-associated proteins), and an elevated rate of constitutive and stimulated vesicular cycling. This 30-50% strengthening of presynapses depends on a set level of actin enrichment, as it disappears after both disassembly or over-stabilization of axonal actin. Acute modulation of actin assembly by Arp2/3 and formins inhibitors results in more subtle effects on presynaptic component concentration and vesicular cycling, suggesting that they target distinct actin structures within presynapses. Capitalizing on the bead-induced presynapse model, we were able to directly visualize these presynaptic actin structures using super-resolution SMLM. Coupled with pharmacological perturbation, SMLM revealed that presynaptic actin forms three main type of nano-structures: an Arp2/3-sensitive actin mesh at the active zone, formin-sensitive actin rails between the active zone and the deeper presynapse containing the reserve pool of synaptic vesicles, and dense, branched actin corrals surrounding the whole presynaptic compartment. This first comprehensive view of presynaptic actin architecture provides a structural basis for multiple previously proposed functions and for new hypotheses on the role of actin at presynapses (Fig. 7E).

Visualizing the arrangement of actin within presynapses presents several challenges. Brain mammalian synapses are usually small and packed with a high density of components, with actin being the most abundant one (Wilhelm et al., 2014). In addition, actin filaments are hard to optimally preserve and not readily visible on classic EM thin sections (Papandréou and Leterrier, 2018). Finally, the close apposition of an even greater amount of postsynaptic actin makes optical fluorescence approaches, even super-resolutive ones, challenging (see Fig. S1). Selective labeling of presynaptic actin using over-expressed probes such as LifeAct-or actin-GFP is possible (Reshetniak and Rizzoli, 2019), but avoiding perturbative effects requires using low labeling densities, below what is necessary to resolve the fine organization of actin filaments. To obtain the highest density of actin labeling using fluorescent phalloidin without interference from the postsynaptic side, we used bead-induced presynapses. This strategy was the first described model of presynaptic induction both in vitro and vivo (Burry, 1982; Burry and Hayes, 1986). How polylysine-coated beads can induce functional presynaptic specialization has long remained unknown (Südhof, 2018), but several more recent studies point to adhesion protein pathways and the recruitment of neuroligin 1 (Lucido et al., 2009; Zhang et al., 2018). We have further validated that bead-induced presynapses concentrate all tested presynaptic components and confirmed that they are competent for synaptic vesicle cycling using anti-synaptotagmin antibody feeding and FM1-43 staining experiments. Finally, we have carefully verified that experiments that do not involve measuring actin content, such as presynaptic component concentration and vesicular cycling after pharmacological treatments, give similar results in bead-induced presynapses and in natural (axon-dendrite) presynapses present in the same neuronal cultures.

Being able to specifically visualize presynaptic actin, we first found that a major subpopulation of presynapses is enriched in actin, and that these actin-enriched presynapses are structurally and functionally stronger than the non-enriched presynapses. Numerous studies have examined the role of actin at presynapses, with a large variability in results and conclusions (Cingolani and Goda, 2008; Papandréou and Leterrier, 2018; Wu and Chan, 2022). Among them, actin disassembly has been shown to impact immature synapses (5-6 div) but not established synapses in mature neurons (12-16 div)(Zhang and Benson, 2001). Here, we have induced presynapses between 8 and 10 div, when neurons are fully competent for presynapse formation. In addition, presynapses form within minutes after bead contact (Suarez et al., 2013), allowing them to maturate over 48 h after seeding in our experiments. We further verified that more mature neurons at 14 div generate the same proportions of actin-enriched and non-enriched presynapses 48 h after seeding, suggesting that these two populations of presynapses are unlikely to be a transient developmental stage of maturation. Actin has also been shown to drive activity-induced unsilencing of presynaptically non-functional synapses in 7-11 div neurons (Shen et al., 2006; Yao et al., 2006). In bead-induced presynapses, we did not see such a clear on/ off difference in vesicular cycling between actin-enriched and non-enriched presynapses. However, absence of actin enrichment is associated with lower presynaptic content, and less constitutive and stimulated cycling. In addition, acutely disassembling actin brings actin-enriched presynapses to the components and cycling levels of non-enriched ones.

What are the presynaptic actin structures that underlie this strengthening effect? Arp2/3 and formins have distinct effects on presynaptic content and vesicular cycling, with further variations depending on the presynaptic component assessed. This pointed to the presence of different actin structure within presynapses, with distinct dependence on Arp2/3 or formins-driven nucleation. Thanks to bead-induced presynapses, we could visualize the nanoscale organization of actin with exquisite resolution. Despite the high variability of shapes and orientation in presynapses at axon-bead contacts, our optimized SMLM consistently revealed the presence of three main actin structures within presynapses: an active zone mesh, rails between the active zone and the deeper reserve pools, and corrals around the whole presynaptic compartment. This direct visualization brings together decades of scattered EM observation and structural speculations from perturbation experiments (Dillon and Goda, 2005), allowing for functional hypotheses. The presence of branched actin at the active zone (Hirokawa et al., 1989) could have a resistive effect on exocytosis (Aunis and Bader, 1988; Morales et al., 2000); the actin rails (Siksou et al., 2007) might help trafficking of vesicles between the reserve pool and the readily-releasable pool at the active zone, or the return of endocytosed vesicles back to the reserve pool (Bloom et al., 2003; Sakaba and Neher, 2003); and the perisynaptic branched actin corrals (Li et al., 2010) could help in constraining, shaping or retaining components of the reserve pool (Sankaranarayanan et al., 2003).

Once these structures have been delineated, it is more straightforward to explain the effects we obtained after actin perturbation on presynaptic component concentration and vesicular cycling. Actin disassembly has a negative effect on components and cycling, but over-stabilization also reduces them due to the strengthening of actin structures resisting synaptic cycling, such as the active zone mesh. Arp2/3 inhibition impacts both the active zone mesh, enhancing release, and the perisynaptic corrals, leading to a reduction of scaffolds but no net change in vesicular cycling. Formins inhibition targets the actin rails, reducing the supply of vesicles from the reserve pool and the overall cycling activity. Of note, the strong effect of formins inhibition is likely partly due to a reduction of inter-synaptic vesicle trafficking by actin trails (Chenouard et al., 2020; Ganguly et al., 2015). It should also be kept in mind that the formins inhibitor SMIFH2 can inhibit non-muscular myosin II (NMII) activity partially. While SMIFH2 is 3-6X less potent on NMII than on formins, NMII is implicated in the synaptic vesicle cycle (Chandrasekar et al., 2013; Peng et al., 2012). However, NMII is differentially implicated in constitutive and evoked release (Peng et al., 2012), a selective effect we do not see when applying SMIFH2 to induced presynapses. Finally, formins implication in the synaptic vesicle cycle has been confirmed independently of SMIFH2 by knockdown of the formin mDia1 (Soykan et al., 2017).

Isolated presynapses in culture are undoubtedly a reductionist approach to the myriad of processes influencing synapse physiology in vivo, but they have allowed us to uniquely isolate and reveal core principles of presynaptic actin organization. Multiple additional layers are added to further shape the presynaptic actin architecture through maturation and plasticity. Feedback mechanisms from the postsynaptic compartments such as cannabinoid regulation of the presynaptic architecture (McFadden et al., 2018) or neuroligin-driven strengthening of presynaptic scaffolds (Wittenmayer et al., 2009) are absent from our experimental model. This also prevents further refinements, like the formation of cross-synaptic nanocolumns that align the release machinery to the clusters of postsynaptic receptors (Tang et al., 2016) – although it should be noted that recent works hints at a primary role of the presynaptic side in organizing nanocolumns (Ramsey et al., 2021). Our study provides the first comprehensive visualization of presynaptic actin structures at the nanoscale, replacing the inferred cartoons of past studies by experimental data. We look forward for this structural insight to help revisiting functional hypotheses and elaborating new ones toward a better understanding of actin multiple roles at presynapses.

## Supporting information

Unformatted manuscript

## Acknowledgments

This work has received support from the CNRS ATIP program (AO 2016 to CL), as well as from from A*MIDEX (ICN PhD Program, ANR-11-IDEX-0001-02 grant) funded by the French Government “Investissements d’Avenir” program (Pépinière to MJP, NeuroSchool PhD program to DB, Master funding to NvB). We would like to thank Subhojit Roy for insightful discussions and Archan Ganguly for the initial help in setting up the bead-induced presynapse model.

## Methods

### Animals, cell culture and polylysine-coated bead treatment

All procedures were in agreements with the guidelines established by the European Animal Care and Use Committee (86/609/CEE) and was approved by the local ethics committee (agreement G13O555). Experiments were performed on pregnant female Wistar rats (Janvier labs). Animals were sacrificed by decapitation and embryo brains were used for primary neuronal cell culture. Rat hippocampal neurons were cultured following the Banker method, above a feeder glia layer (Kaech and Banker, 2006). Rapidly, 18-mm diameter round, #1.5H coverslips were affixed with paraffine dots as spacers, then treated with poly-L-lysine. Hippocampi from E18 rat pups were dissected, and homogenized by trypsin treatment followed by mechanical trituration and seeded on the coverslips at a density of 6,000 cells/cm2 for 3 hours in serum-containing plating medium (MEM with 10% fetal bovine serum, 0.6% added glucose, 0.08 mg/mL sodium pyruvate, 100 UI/mL penicillin-streptomycin). Coverslips were then transferred, cells down, to petri dishes containing confluent glia cultures conditioned in NB+ medium (Neurobasal medium supplemented with 2 % B-27, 100 UI/mL penicillin/ streptomycin and 2.5 µg/mL amphotericin) and cultured in these dishes for up to 3 weeks. For most bead-induced presynapse experiments (except Fig. S3), neurons were cultured for 8 days in vitro (8 div) before the addition of beads. Poly-D-lysine beads are prepared as follows: 30 million 4.5 µm aliphatic amine latex beads (Thermo Fisher) were incubated with poly-D-lysine (P7405, Sigma, 63 µg/mL) in Dulbecco’s phosphate buffered saline (PBS, Thermo Fisher) for 3 h at room temperature with rotary agitation. Beads were then washed twice in sterile water and once with NB+ medium. Beads were resuspended in NB+ medium and added dropwise to the neurons at a concentration of 3 million beads/coverslip, neurons facing up. After 3 h, the coverslips were then flipped back, neurons facing the astrocytes for two days before treatments.

### Pharmacological treatments and anti-synaptotagmin vesicular cycling assays

2 days after bead seeding (10 div), neurons were treated 1h with 1 µM swinholide A (Sigma), 3 h with 5 µM cucurbitacin E (Sigma), 1 h with 50 µM CK666 (Bio-Techne) or 1 h with 30 µM SMIFH2 (Bio-Techne). Drugs were directly added in the culture medium and controls included the same amount of dimethyl sulfoxide (DMSO) that was originally used to dissolve the drugs. Neurons were rinsed 3 times with PBS prior to fixation.

Vesicular cycling assays were performed using an antibody directed to an extracellular epitope of synaptotagmin 1 (syt, clone 604.2, Synaptic Systems) that was fed to neurons 2 days after bead seeding (10 div). For assaying constitutive cycling, neurons were incubated with the syt antibody at 1.6 µg/µL in HBS medium (NaCl 136 mM, HEPES 10 mM, D-glucose 10 mM, CaCl2 2 mM, MgCl2 1,3 mM) for 1 h at 37°C, 5% CO2. For assaying stimulated cycling, neurons were treated with the syt antibody in HBS medium supplemented with 50 mM KCl for 3 min at room temperature. Cells were then washed 3 times with PBS before fixation.

### FM1-43 experiments on living cells

2 days after seeding with polylysine-coated beads, neurons were incubated for 1 h at 37 °C, 5% CO2 in NB+ medium containing 100 nM SiR-actin (Spirochrome). Neurons were then incubated in HBS medium for 10 minutes at 37 °C. Following this incubation, the neurons were transferred to live-cell imaging chamber (Life Imaging Services) containing HBS medium and placed at 37°C in the incubated chamber of an epifluorescence/TIRF microscope (Zeiss). HBS medium was then exchanged for HBS/KCL medium containing APV at 50 µM, CNQX at 25 µM and FM1-43 (SynaptoGreen-C4, Biotium) at 5 µM for 180 seconds at 37 ºC. HBS/KCL medium was then changed back to HBS medium after 3 times 1 min 30 s rinses with HBS medium. Neurons were then left to recover for 15 minutes in HBS medium. Green (FM1-43) and far-red (SiR-actin) channel epifluorescence images were taken using a 40X objective for several fields of view selected for the presence of a number of bead-induced. FM1-43 release was achieved following an additional 180 s incubation with HBS/KCl medium with APV/CNQX, followed by 3 rinses with HBS medium every 1 min 30 s, and 15 min recovery in HBS medium. Post-release images of the same preselected fields of view were then acquired using the same settings as the post-loading images.

### Immunocytochemistry and sample preparation

2 days after bead seeding (10 div), neurons were fixed and stained similarly to published procedures (Jimenez et al., 2020). Fixation used 4% PFA in PEM buffer (PIPES 80 mM, MgCl2 2 mM, EGTA 5 mM, pH 6.8) for 10 min at room temperature. After rinses in 0.1 M phosphate buffer (TpO4), neurons were blocked for 1 h in immunochemistry buffer (ICC: TpO4, gelatin 0.22%, Triton 0.1%) and incubated with primary antibodies diluted in ICC, overnight at 4°C. Primary antibodies used are chicken anti-map2 (a5392, 1:1000, Abcam), mouse anti-ß2-spectrin (#612563, 1:150, BD Bioscience), guinea pig anti-synaptophysin (guinea pig, 101 004, 1:400, Synaptic Systems), rabbit anti-synapsin (ab1543P, 1:1000, Merck), mouse anti-bassoon (clone SAP7F407, 1:200, Abcam), mouse anti-vamp2 (clone 69.1, 1:200, Synaptic Systems), guinea pig anti-VGLUT (135 304, 1:500, Synaptics Systems) and rabbit anti-VGAT (131 003, 1:1000, Synaptic Systems). After rinses in ICC, corresponding secondary antibodies conjugated to Alexa Fluor 488, 555 or 647 (Thermo Fisher) or DyLight 405 (Rockland) diluted in ICC (1:200 to 1:400) were incubated for 1 h at room temperature. Secondary antibodies were rinsed, and neurons were incubated with Alexa Fluor 647-conjuguated phalloidin (A2287, 1:40 in TpO4, ThermoFisher) or Atto 488-conjugated phalloidin (#AD488-81, 1:40, Atto-Tec) for 1h30 at room temperature or overnight at 4°C. For epifluorescence imaging, coverslips were mounted in Prolong Glass containing fluorescent phalloidin (1:400). Alternatively, after the rinsing steps, stained coverslips were kept in PB + 0.02% sodium azide and Alexa Fluor 647 phalloidin (1:40) at 4°C before STORM imaging.

### Epifluorescence microscopy

Diffraction-limited epifluorescence images were obtained using an Axio-Observer upright microscope (Zeiss) equipped with a 40X NA 1.4 or 63X NA 1.4 objective and an Orca-Flash4.0 camera (Hamamatsu). Appropriate hard-coated filters and dichroic mirrors were used for each fluorophore. A thin Z-stack of 3-10 slices spaced by 0.2 µm was acquired to include the signal from all neuronal processes within the whole field of view. For illustration images, image editing was performed using Fiji (Schindelin et al., 2012) and included projection using the Extended Depth of Field plugin (Forster et al., 2004), linear contrast adjustment, and gamma adjustment (0.7-0.9) to highlight the faint actin labeling along axons. In the case of different treatment conditions labeled for the same targets, the contrast settings were kept constant for each channel across the different conditions/treatments.

### Single Molecule Localization Microscopy

For single color SMLM, we used Stochastic Optical Microscopy (STORM). STORM was performed on an N-STORM microscope (Nikon Instruments). Stained coverslips were mounted in a silicone chamber filled with STORM buffer (Smart Buffer Kit, Abbelight). The N-STORM system uses an Agilent MLC-400B laser launch with 405 nm (50 mW maximum fiber output power), 488 nm (80 mW), 561 mW (80 mW) and 647 nm (125 mW) solid-state lasers, a 100X NA 1.49 objective and an Ixon DU-897 camera (Andor). After locating an area with bead-synapses using low-intensity illumination, a TIRF image was acquired, followed by a STORM acquisition. 30,000-60,000 images (256×256 pixels, 15 ms exposure time) were acquired at full 647 nm laser power. Reactivation of fluorophores was performed during acquisition by increasing illumination with the 405 nm laser. When imaging actin, 30 nM phalloidin-Alexa Fluor 647 was added to the STORM buffer to mitigate actin unbinding during imaging (Jimenez et al., 2020). For three-color SMLM (Fig. 6F), we used STORM in combination with DNA-PAINT (Jimenez et al., 2020). Neurons were labeled using rabbit anti-synapsin and mouse anti-ß2-spectrin primary antibodies (see above), then anti-rabbit and anti-mouse secondary antibodies coupled to distinct PAINT DNA handles, as well as phalloidin-Alexa Fluor 647 for actin. Imaging was done sequentially, first for actin in STORM buffer (60,000 frames at 67 Hz), then for two immunostained proteins in PAINT buffer (0.1M phosphate buffer saline, 500 mM NaCl, 5% dextran sulfate, pH 7.2) supplemented with 0.12-0.25 nM of PAINT imagers strands coupled to Atto565 and Atto650 (Metabion, 2 times 40,000 frames of alternating 561 nm and 647 nm excitation at 30 Hz).

For 2D STORM and PAINT images (Fig. S1 and Fig. 6), the N-STORM software (Nikon Instruments) was used for the localization of single fluorophore activations, and correct for drift using cross-correlation. For two-color PAINT acquisitions using alternating 561 nm and 647 nm excitation, lateral chromatic aberration correction was corrected within the N-STORM software using polynomial warping calibrated on sub-diffraction beads. The list of localizations was then exported as a text file. For 3D STORM images (Fig. S1 and Fig. 7), acquired stacks were processed using DECODE (Speiser et al., 2021). Briefly, PSF were modeled using spline fitting in SMAP (Li et al., 2018) and used to simulate sequences of blinking events using characteristics (photon number range, lifetime distribution) inferred from real acquisition data from the N-STORM microscope. A Pytorch model was trained to infer the 3D coordinates and uncertainty of the simulated blinking events, then applied to the experimental acquired sequences (Speiser et al., 2021). The resulting localizations (fitted blinking events) were filtered based on uncertainty, and drift during acquisition was corrected in 3D using a redundant cross-correlation algorithm (Wang et al., 2014) implemented as an independent module of SMAP (Ries, 2020). After translation of the coordinate files obtained from N-STORM or DECODE/SMAP, image reconstructions were performed using the ThunderSTORM ImageJ plugin (Ovesny et al., 2014) in Fiji software. Custom scripts and macros were used to translate coordinate files, as well as automate images reconstruction for whole images at 16 nm/pixel and detailed zooms at 4 nm/pixel (https://github.com/cleterrier/ChriSTORM). For visualization of faint actin structures, the local contrast of monochrome (2D) or Z-colored (3D) reconstructed images was enhanced using the Contrast Limited Adaptive Histogram Equalization (CLAHE) plugin.

### Epifluorescence image analysis

Intensity quantifications were performed on maximum projection of the raw data with no further adjustment (see below). Linear regions of interests (ROIs) were traced along an axon using the NeuronJ plugin in Fiji software, encompassing an axon-bead contact. Tracings were then translated into ImageJ ROIs. The ROI was then refined to the presynaptic cluster using the ProFitFeat script (available at https://github.com/cleterrier/Measure_ROIs/blob/master/Pro_Feat_Fit.js), restricting the ROI to the segment with an intensity above 50% of the maximum intensity point along the line ROI. The background-corrected intensities within these ROIs were then measured for each labeled channel. For natural synapses, ROIs were traced along the axon at axon-dendrite contacts where presynaptic proteins were visibly accumulated and fluorescence intensities were measured for each labeled channel as described above.

### STORM image analysis

Processed STORM localization files were used to obtain high-magnification images (4 nm/pixel) of each bead-induced presynapse imaged from 2 to 5 independent experiments (6 for control, 32 for CK66 treatment, 18 for SMIFH2 treatment). Actin structures present on the reconstructed image were scored as present, non-present, or not determined when their presence or absence could not be reliably determined.

### Data presentation and statistical analysis

Sup. File 1 recapitulates the statistics for all quantifications shown as graphs in the Figures. Individual measurement points (number n) from independent experiments (number N) were pooled. All experiments were replicated between 2-and 8-times using bead-seeded neurons from different cultures. Intensity profiles, graphs and statistical analyses were generated using Prism. On bar graphs, dots (if present) are averages of each independent experiment, bars or horizontal lines represent the mean, and vertical lines are the SEM unless otherwise specified. Significances were tested using one-way, non-parametric ANOVA with Šídák post-hoc significance testing between selected conditions. All differences and variations mentioned in the text are significant, unless specified as non-significant. In Sup. File 1, the results of the post-hoc significance are indicated as: ns non-significant * p < 0.05, ** p < 0.01, *** p < 0.001.

---

**Figure S1:**
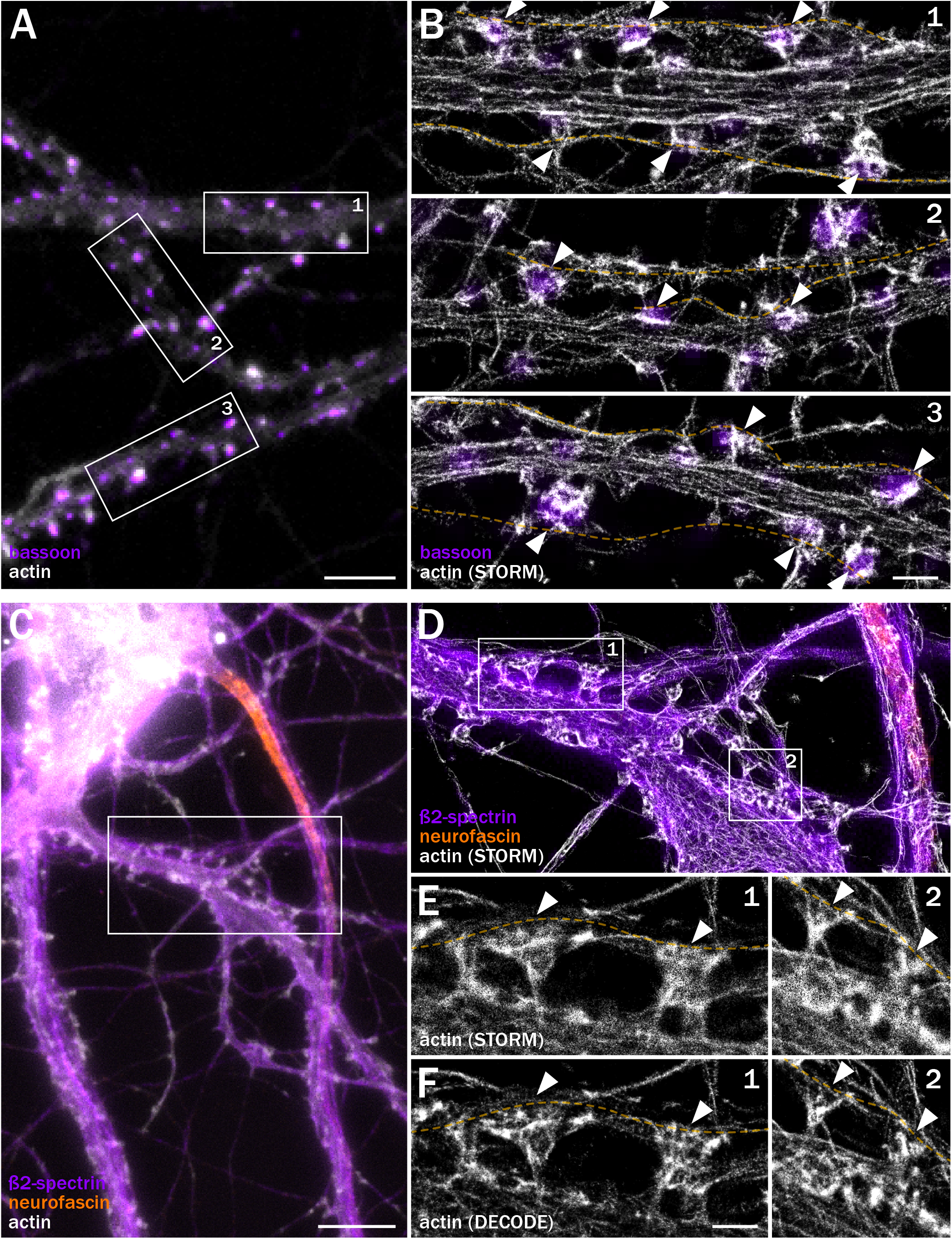
Postsynaptic actin impedes the visualization of presynaptic content and organization in hippocampal neurons, even using state-of-the-art super-resolution microscopy. A-B. Images of neurons after 20 div, stained for bassoon (purple) and actin (gray). A, widefield image. B, zooms from the highlighted areas in A, overlaying the widefield image of bassoon with the STORM image of actin, showing the actin-enriched dendritic spines (arrowheads) that mask the fainter presynaptic actin along axons (yellow dashed lines). C-F. Images of neurons after 20 div, stained for ß2-spectrin (purple), neurofascin (orange) and actin (gray). C, widefield image. D, zoom from the highlighted areas in C, overlaying widefield images of ß2-spectrin and neurofascin with the STORM image of actin. E, isolated STORM image of actin further zoomed from the areas highlighted in D, showing the actin-enriched dendritic spines (arrowheads) that mask the fainter presynaptic actin along axons (yellow dashed lines). E, Enhanced actin STORM image of the same areas obtained using deep learning-based DECODE processing, allowing for a better definition of bright actin structures in dendritic spines (arrowheads). Postsynaptic actin still impedes the clear visualization of presynaptic actin along axons (yellow dashed lines). Scale bars for A and C, 10 µm; for B, D and E, 1 µm.

**Figure S2:**
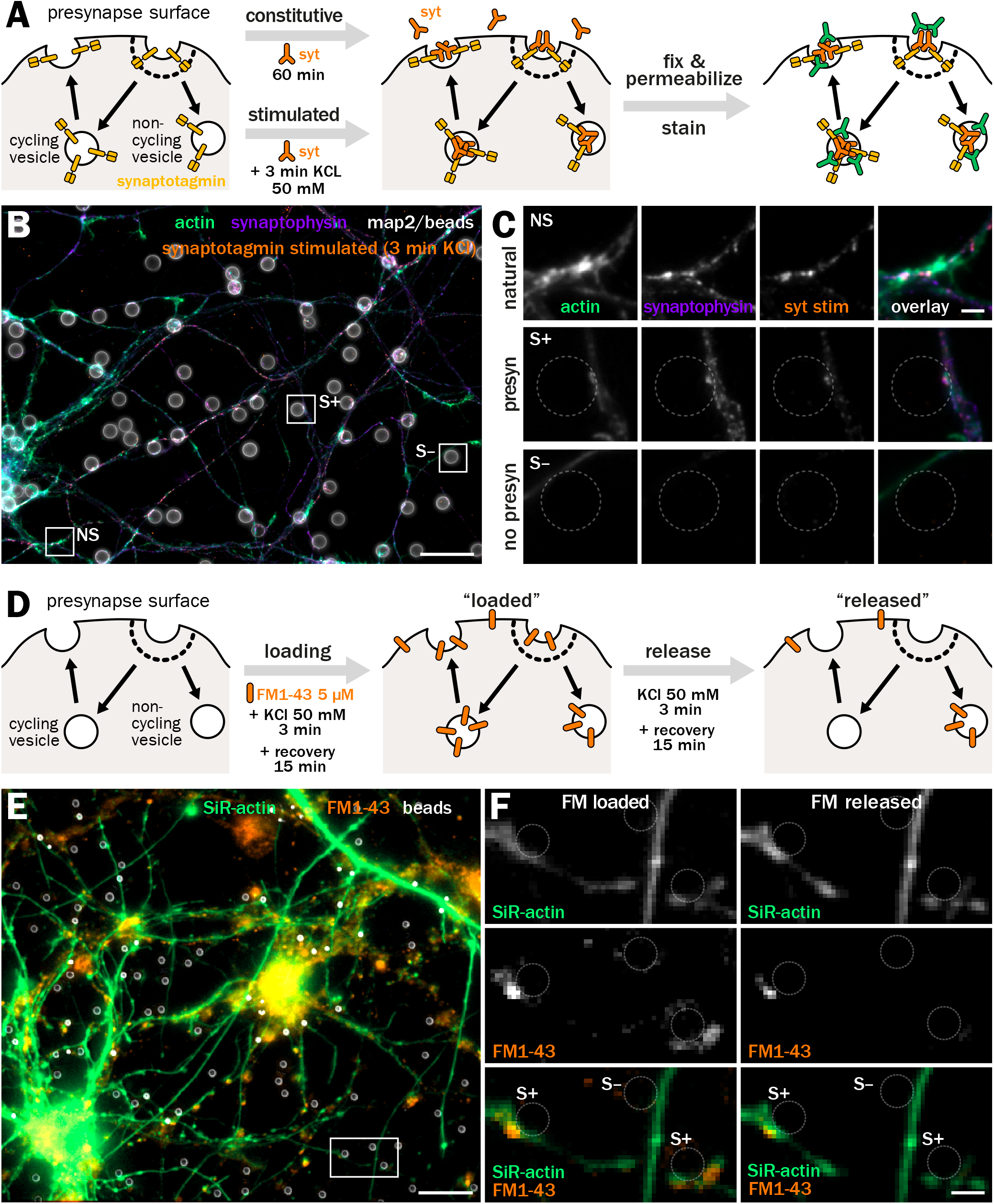
Optical methods to assess vesicular cycling at bead-induced presynapses. A-C. Anti-synaptotagmin (syt) antibody feeding experiments to assess the cycling activity of presynapses. A, cartoon of the anti-syt feeding experiments. Living neurons are incubated with the syt antibody (orange) directed against an extracellular epitope of synaptotagmin (yellow), either for 60 minutes to measure constitutive cycling, or for 3 minutes in the presence of 50 mM KCl to measure stimulated cycling. Neurons are then fixed and the syt antibody is revealed with a secondary antibody (green). B, widefield fluorescence image of cultured neurons 2 days after bead seeding at 8 div, labeled for actin (green), synaptophysin (purple), map2 (gray), and feeding with anti-synaptotagmin antibody (syt) during a 3 min incubation with KCl (stimulated cycling). C, zooms corresponding to the natural synapse (NS), S+ and S– axon-bead contacts highlighted in B. Scale bars for B, 20 µm; for C, 2 µm. D-E. FM1-43 dye loading/release experiment to assess the cycling activity of presynapses. D, cartoon of the FM1-43 experiment. Living neurons are first loaded with FM1-43 using a 3 min incubation in 50 mM KCl followed by 15 min of recovery, then images of the “loaded” time point are taken. FM1-43 is then released using a second 3-min incubation with 50 mM KCl and 15-min recovery, before images are taken of the “released” time point. E, widefield fluorescence image of cultured neurons 2 days after bead seeding at 8 div, labeled for actin using SiR (green) after loading with FM1-43 (orange). F, zooms corresponding to area highlighted in E, showing the images obtained after loading (left column) and release (right column) of FM1-43. Beads are indicated by dashed-line circles labeling the induced presynapses (S+) and axon-bead contact with no presynapse (S–). Scale bars for B, 20 µm; for C, 5 µm.

**Figure S3:**
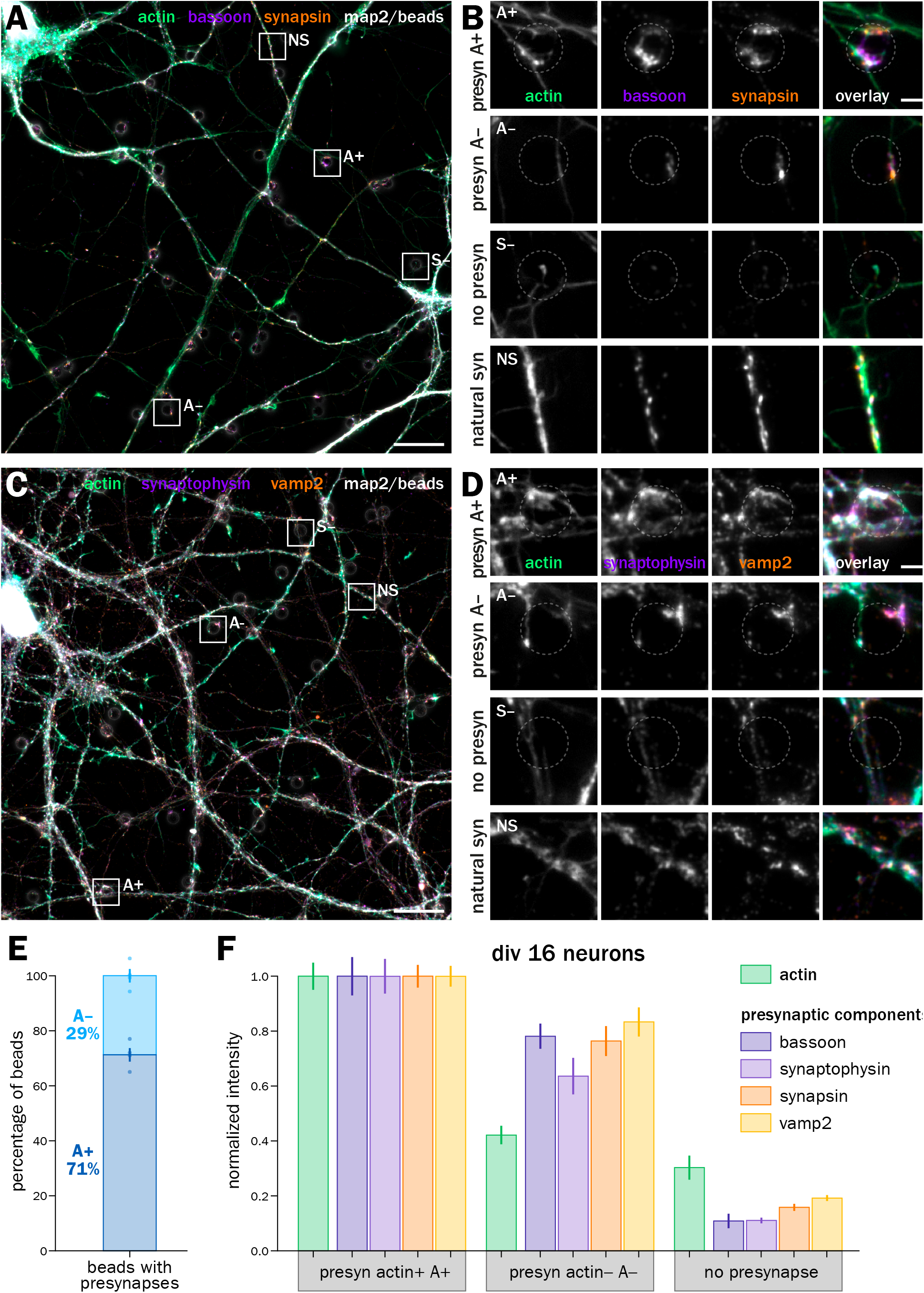
Presynapses induced on 14-div neurons show a similar proportion of actin enrichment and with similar presynaptic content. A. Widefield fluorescence image of cultured neurons 2 days after bead seeding at 14 div, labeled for actin (green), bassoon (purple), synapsin (orange), and map2 (gray). B. Zooms corresponding to the areas highlighted in A: top row, actin-enriched induced presynapse at an axon-bead contact (A+); second row, induced presynapse at an axon-bead contact with no actin enrichment (A–); third row, axon-bead contact with no induced presynapse (S–); bottom row, natural synapse at axon-dendrite contact (NS). C. Widefield fluorescence image of cultured neurons 2 days after bead seeding at 14 div, labeled for actin (green), synaptophysin (purple), vamp2 (orange), and map2 (gray). D. Zooms corresponding to the A+, A–, S– axon-bead contacts and natural synapses (NS) highlighted in C. Scale bars in A, C: 20 µm; B, D: 2 µm. E. Quantification of the proportion of A+ (dark blue) and A– (light blue) at axon-bead contacts that resulted in an induced presynapse. F. Quantification of the labeling intensity for actin (green), bassoon (dark purple), synaptophysin (purple), synapsin (orange), and vamp2 (yellow) at actin-enriched presynapses (A+), induced presynapses with no actin enrichment (A–), and axon-bead contacts devoid of presynapse (S–), normalized to the intensity at A+ presynapses.

**Figure S4:**
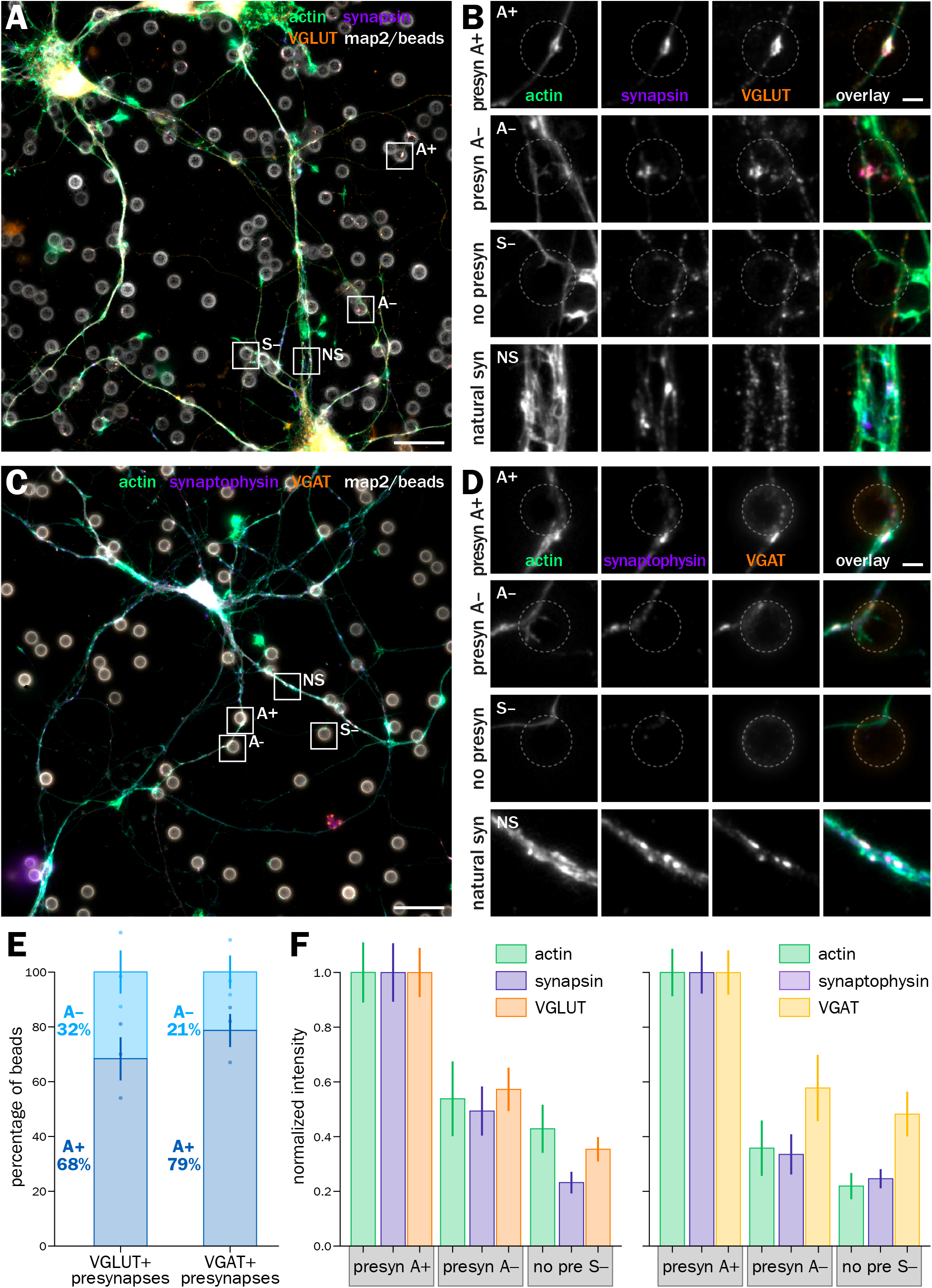
Bead-induced inhibitory and excitatory presynapses show similar actin and presynaptic component content distribution. A. Widefield fluorescence image of cultured neurons 2 days after bead seeding at 8 div, labeled for actin (green), synapsin (purple), VGLUT (orange), and map2 (gray). B. Zooms corresponding to the areas highlighted in A: top row, actin-enriched induced presynapse at an axon-bead contact (A+); second row, induced presynapse at an axon-bead contact with no actin enrichment (A–); third row, axon-bead contact with no induced presynapse (S–); bottom row, natural synapse at axon-dendrite contact (NS). C. Widefield fluorescence image of cultured neurons 2 days after bead seeding at 8 div, labeled for actin (green), synaptophysin (purple), VGAT (orange), and map2 (gray). D. Zooms corresponding to the A+, A–, S– axon-bead contacts and natural synapses (NS) highlighted in C. Scale bars in A, C: 20 µm; B, D: 2 µm. E. Quantification of the proportion of A+ (dark blue) and A– (light blue) at axon-bead contacts that resulted in an induced VGLUT-positive (left bar) and VGAT-positive (right bar) presynapse. F. Left graph, quantification of the labeling intensity for actin (green), synapsin (dark purple) and VGLUT (orange) at actin-enriched presynapses (A+), induced presynapses with no actin enrichment (A–), and axon-bead contacts devoid of presynapse (S–), normalized to the intensity at A+ presynapses. Right graph, quantification of the labeling intensity for actin (green), synaptophysin (purple), and VGAT (yellow) at A+ and A– induced presynapses as well as S– axon bead contacts.

**Figure S5:**
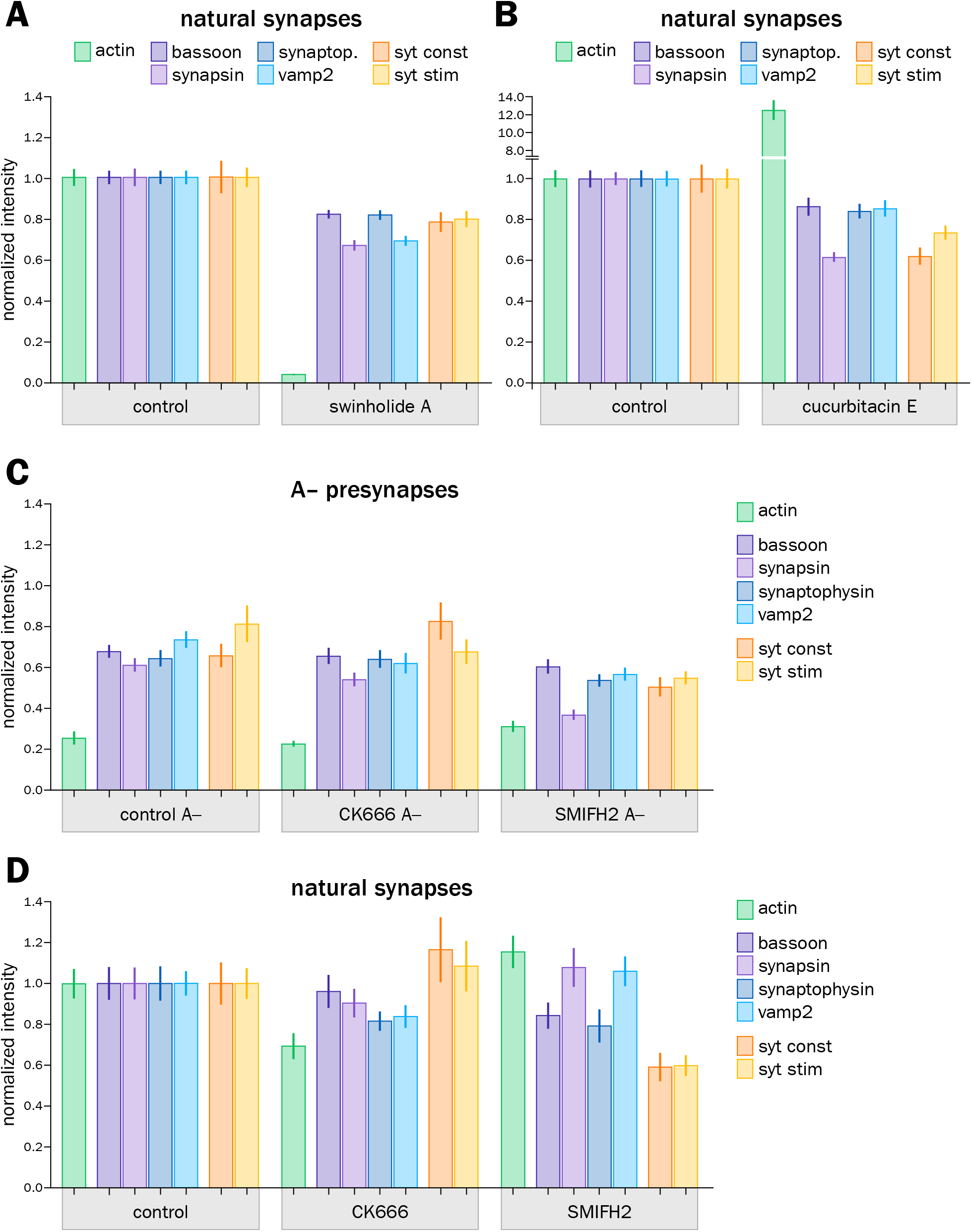
Acute actin perturbation effect on the concentration of presynaptic components is similar at natural synapses and induced presynapses. A. Quantification of the labeling intensity for actin (green), bassoon (dark purple), synapsin (purple), synaptophysin (dark blue), vamp2 (blue), and after syt feeding for constitutive (orange) and stimulated (yellow) vesicular cycling at natural synapses in the control condition and after swinholide A treatment. This was performed on the same images as the induced presynapses quantified in Fig. 4D. B. Quantification of the labeling intensity for actin (green), bassoon (dark purple), synapsin (purple), synaptophysin (dark blue), vamp2 (blue), and after syt feeding for constitutive (orange) and stimulated (yellow) vesicular cycling at natural synapses in the control condition and after cucurbitacin E treatment. This was performed on the same images as the induced presynapses quantified in Fig. 4E. C. Quantification of the labeling intensity for actin (green), bassoon (dark purple), synapsin (purple), synaptophysin (dark blue), vamp2 (blue), and after syt feeding for constitutive (orange) and stimulated (yellow) vesicular cycling at induced presynapses devoid of actin enrichment (A–) in the control condition and after CK666 or SMIFH2 treatment. This was performed on the same images as the A+ induced presynapses quantified in Fig. 5D. D. Quantification of the labeling intensity for actin (green), bassoon (dark purple), synapsin (purple), synaptophysin (dark blue), vamp2 (blue), and after syt feeding for constitutive (orange) and stimulated (yellow) vesicular cycling at natural synapses in the control condition and after CK666 or SMIFH2 treatment. This was performed on the same images as the A+ induced presynapses quantified in Fig. 5D.

