## Supplementary material for "Distinct nano-structures support a multifunctional role of actin at presynapses": Unformatted manuscript

***Figure S5: Acute actin perturbation effect on the concentration of presynaptic components is similar at natural synapses and induced presynapses.***

**A.** Quantification of the labeling intensity for actin (green), bassoon (dark purple), synapsin (purple), synaptophysin (dark blue), vamp2 (blue), and after syt feeding for constitutive (orange) and stimulated (yellow) vesicular cycling at natural synapses in the control condition and after swinholide A treatment. This was performed on the same images as the induced presynapses quantified in Fig. 4D. **B.** Quantification of the labeling intensity for actin (green), bassoon (dark purple), synapsin (purple), synaptophysin (dark blue), vamp2 (blue), and after syt feeding for constitutive (orange) and stimulated (yellow) vesicular cycling at natural synapses in the control condition and after cucurbitacin E treatment. This was performed on the same images as the induced presynapses quantified in Fig. 4E. **C.** Quantification of the labeling intensity for actin (green), bassoon (dark purple), synapsin (purple), synaptophysin (dark blue), vamp2 (blue), and after syt feeding for constitutive (orange) and stimulated (yellow) vesicular cycling at induced presynapses devoid of actin enrichment (A–) in the control condition and after CK666 or SMIFH2 treatment. This was performed on the same images as the A+ induced presynapses quantified in Fig. 5D. **D.** Quantification of the labeling intensity for actin (green), bassoon (dark purple), synapsin (purple), synaptophysin (dark blue), vamp2 (blue), and after syt feeding for constitutive (orange) and stimulated (yellow) vesicular cycling at natural synapses in the control condition and after CK666 or SMIFH2 treatment. This was performed on the same images as the A+ induced presynapses quantified in Fig. 5D.

### Figure 1

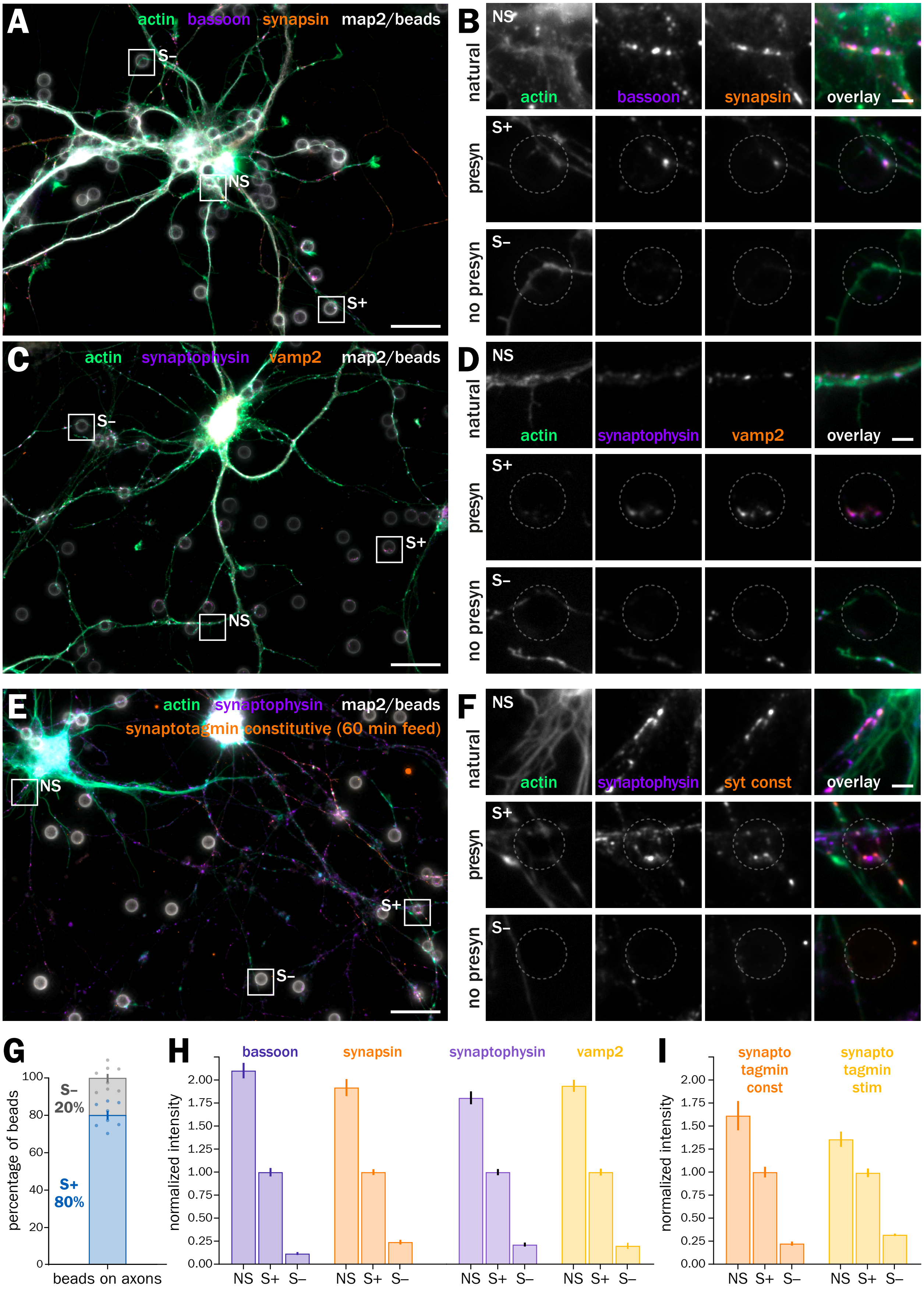
A
actin bassoon synapsin map2/beads
S–
NS
S+
B
NS
natural
actin
bassoon
synapsin
overlay
S+
presyn
S–
no presyn
C
actin synaptophysin vamp2 map2/beads
S–
S+
NS
D
NS
natural
actin
synaptophysin
vamp2
overlay
S+
presyn
S–
no presyn
E
actin synaptophysin map2/beads
synaptotagmin constitutive (60 min feed)
NS
S+
S–
F
NS
natural
actin
synaptophysin
syt const
overlay
S+
presyn
S–
no presyn
G
percentage of beads
S– 20%
S+ 80%
beads on axons
H
normalized intensity
bassoon
synapsin
synaptophysin
vamp2
NS S+ S–
I
normalized intensity
synapto tagmin const
synapto tagmin stim
NS S+ S–

### Figure 2

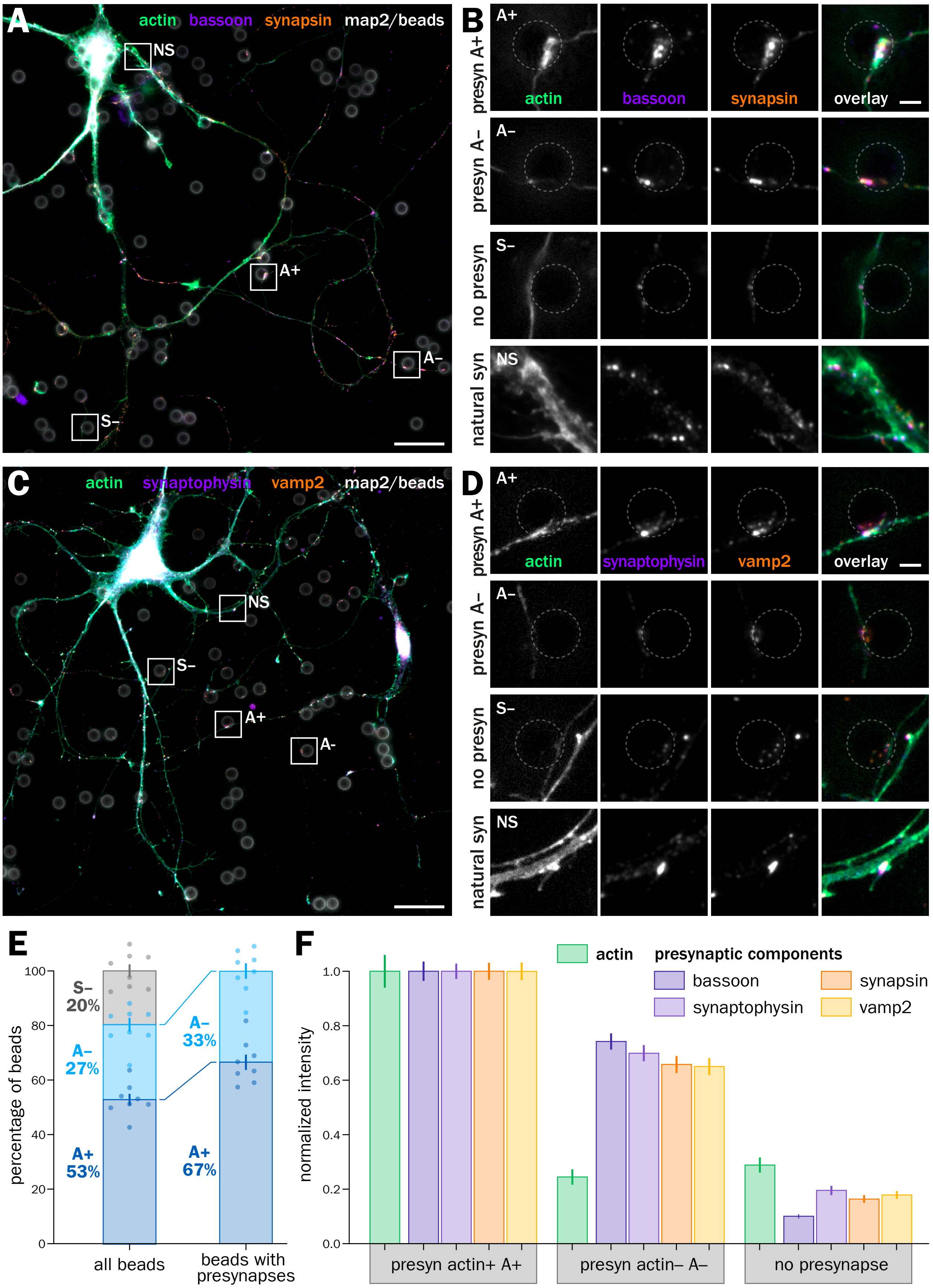

A
actin bassoon synapsin map2/beads
NS
A+
A–
S–
B
presyn A+
presyn A–
no presyn
natural syn
A+
A–
S–
NS
actin
bassoon
synapsin
overlay
C
actin synaptophysin vamp2 map2/beads
NS
S–
A+
A-
D
presyn A+
presyn A–
no presyn
natural syn
actin
synaptophysin
vamp2
overlay
E
percentage of beads
0
20
40
60
80
100
S– 20%
A– 27%
A+ 53%
A– 33%
A+ 67%
all beads
beads with presynapses
F
normalized intensity
0.0
0.2
0.4
0.6
0.8
1.0
actin
presynaptic components
bassoon
synaptophysin
synapsin
vamp2
presyn actin+ A+
presyn actin– A–
no presynapse

### Figure 3

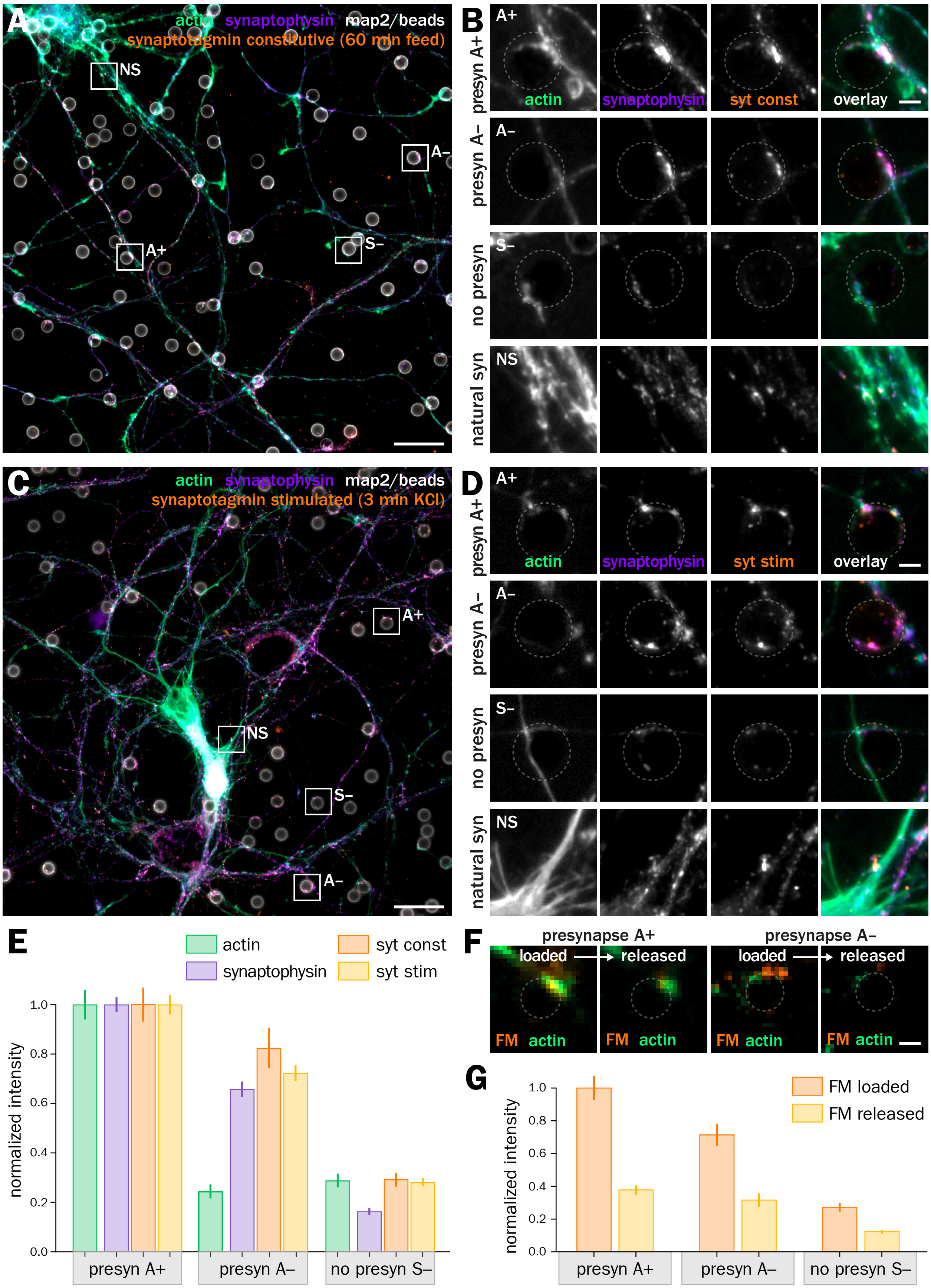
A
actin synaptophysin map2/beads
synaptotagmin constitutive (60 min feed)
NS
A–
A+
S–
B
A+
presyn A+
actin
synaptophysin
syt const
overlay
A–
presyn A–
S–
no presyn
NS
natural syn
C
actin synaptophysin map2/beads
synaptotagmin stimulated (3 min KCl)
D
syt stim
E
actin
synaptophysin
syt const
syt stim
normalized intensity
1.0
0.8
0.6
0.4
0.2
0.0
presyn A+
presyn A–
no presyn S–
F
presynapse A+
presynapse A–
loaded
released
FM actin
G
FM loaded
FM released

### Figure 4

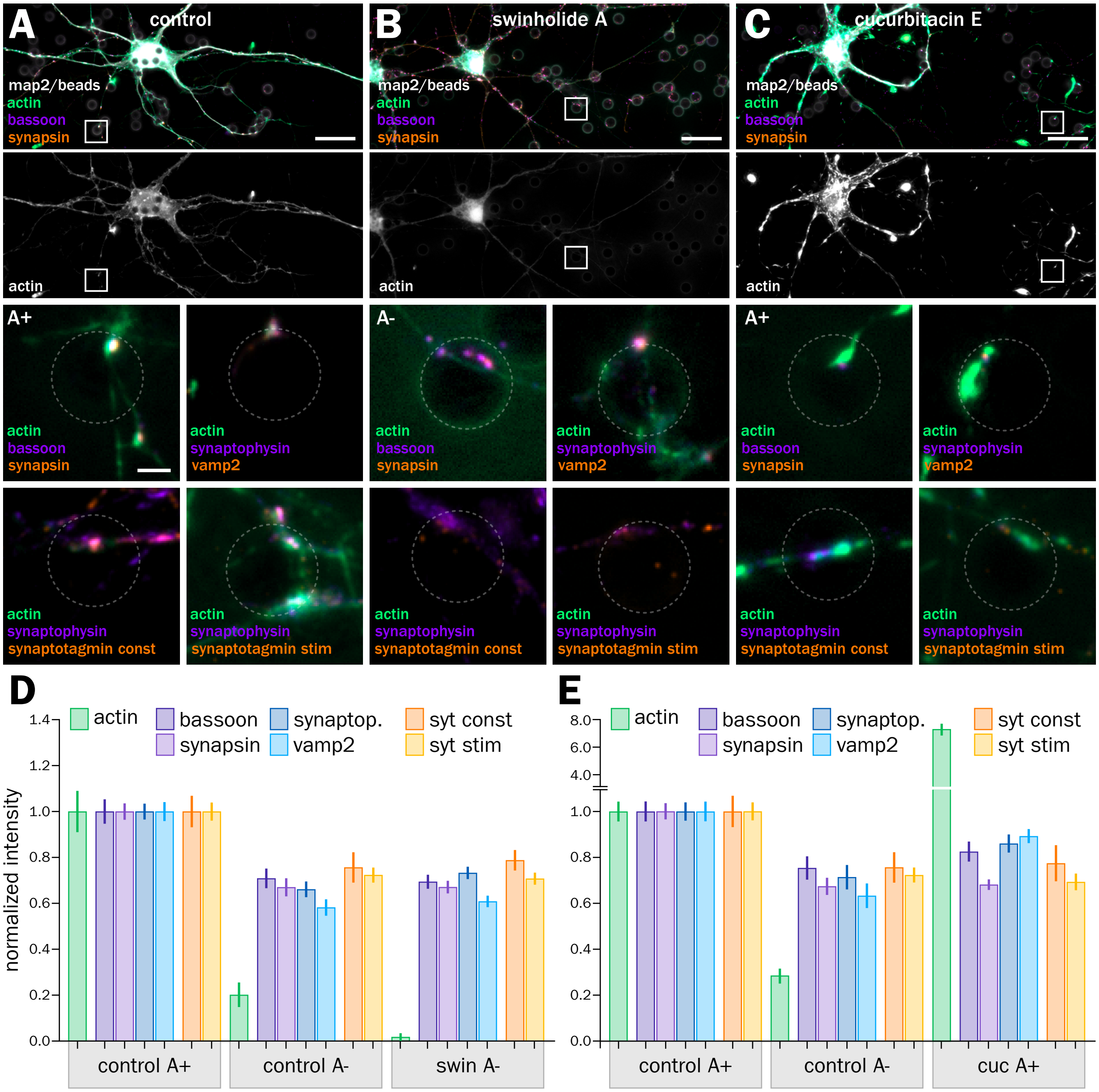
A
control
map2/beads
actin
bassoon
synapsin
actin
B
swinholide A
map2/beads
actin
bassoon
synapsin
actin
C
cucurbitacin E
map2/beads
actin
bassoon
synapsin
actin
A+
actin
bassoon
synapsin
actin
synaptophysin
vamp2
A-
actin
bassoon
synapsin
actin
synaptophysin
vamp2
A+
actin
bassoon
synapsin
actin
synaptophysin
vamp2
actin
synaptophysin
synaptotagmin const
actin
synaptophysin
synaptotagmin stim
actin
synaptophysin
synaptotagmin const
actin
synaptophysin
synaptotagmin stim
actin
synaptophysin
synaptotagmin const
actin
synaptophysin
synaptotagmin stim
D
actin
bassoon
synapsin
synaptop.
vamp2
syt const
syt stim
normalized intensity
1.4
1.2
1.0
0.8
0.6
0.4
0.2
0.0
control A+
control A-
swin A-
E
actin
bassoon
synapsin
synaptop.
vamp2
syt const
syt stim
8.0
6.0
4.0
1.0
0.8
0.6
0.4
0.2
0.0
control A+
control A-
cuc A+

### Figure 5

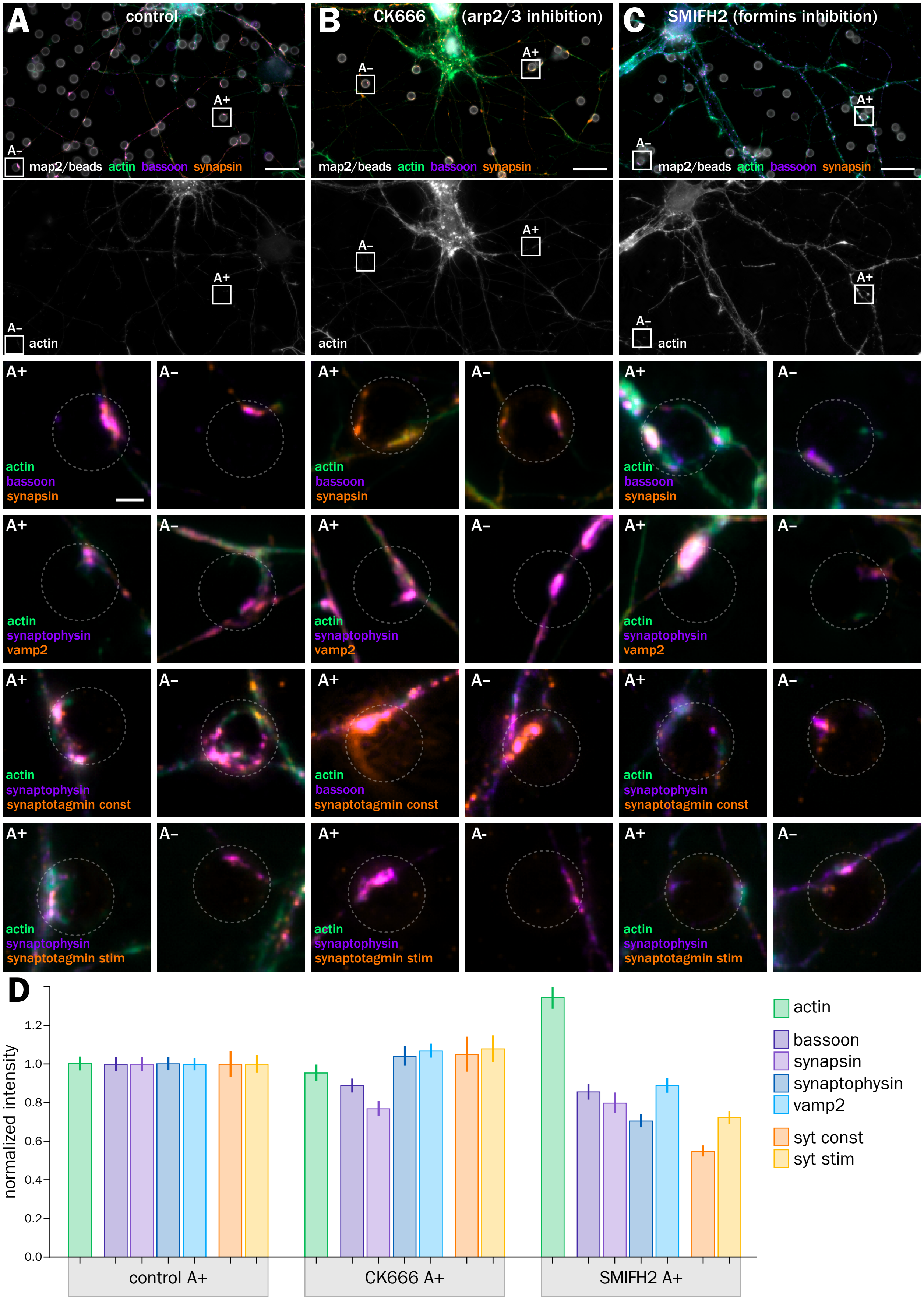
A
control
B
CK666
(arp2/3 inhibition)
C
SMIFH2 (formins inhibition)
A+
A–
map2/beads actin bassoon synapsin
actin
actin
bassoon
synapsin
synaptophysin
vamp2
synaptotagmin const
synaptotagmin stim
D
normalized intensity
0.0
0.2
0.4
0.6
0.8
1.0
1.2
control A+
CK666 A+
SMIFH2 A+
actin
bassoon
synapsin
synaptophysin
vamp2
syt const
syt stim

### Figure 6

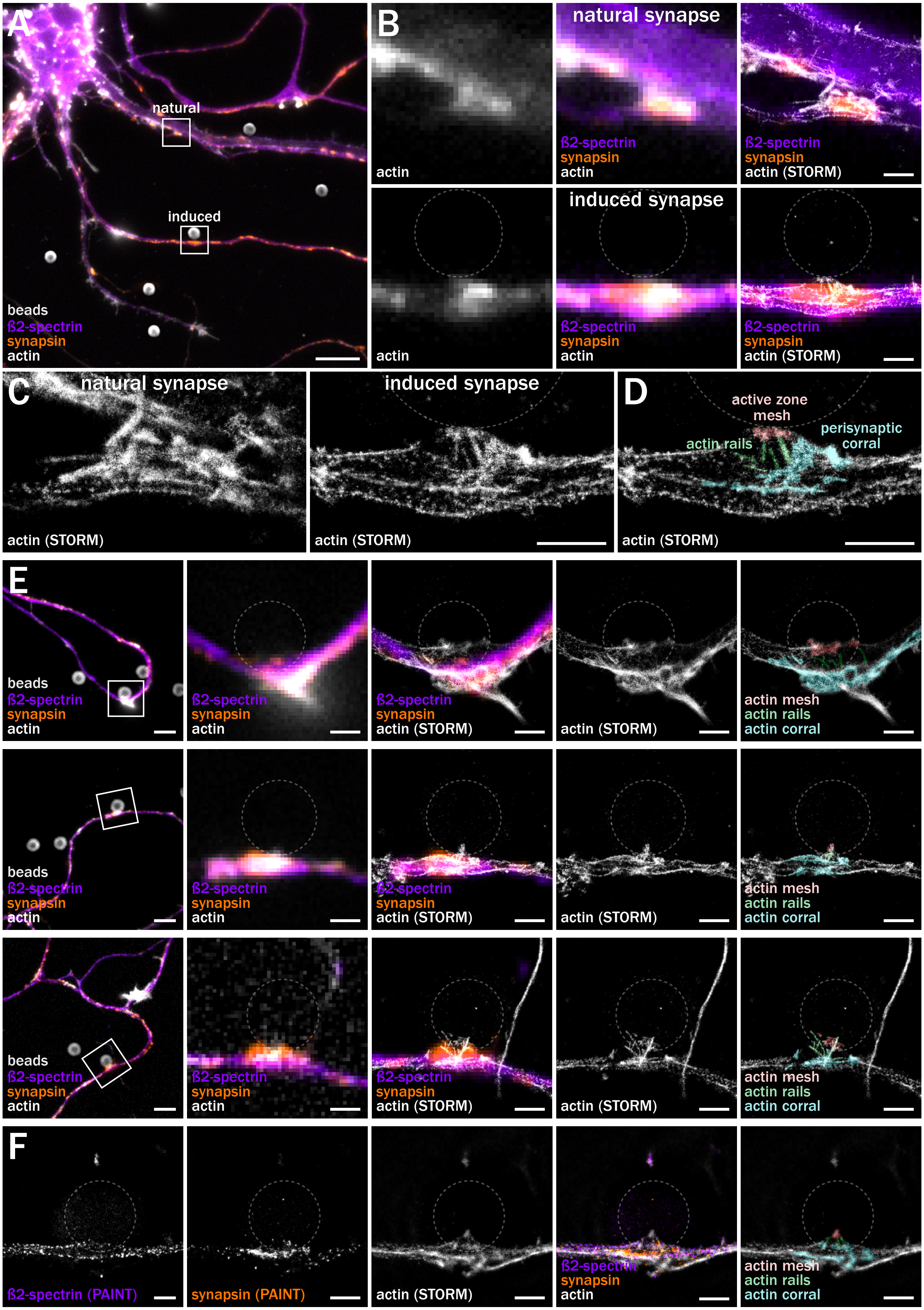
A
natural
induced
beads
ß2-spectrin
synapsin
actin
B
actin
natural synapse
ß2-spectrin
synapsin
actin
ß2-spectrin
synapsin
actin (STORM)
actin
induced synapse
ß2-spectrin
synapsin
actin
ß2-spectrin
synapsin
actin (STORM)
C
natural synapse
actin (STORM)
induced synapse
actin (STORM)
D
active zone
mesh
perisynaptic
corral
actin rails
actin (STORM)
E
beads
ß2-spectrin
synapsin
actin
ß2-spectrin
synapsin
actin (STORM)
actin (STORM)
actin mesh
actin rails
actin corral
F
ß2-spectrin (PAINT)
synapsin (PAINT)
actin (STORM)
ß2-spectrin
synapsin
actin
actin mesh
actin rails
actin corral

### Figure 7

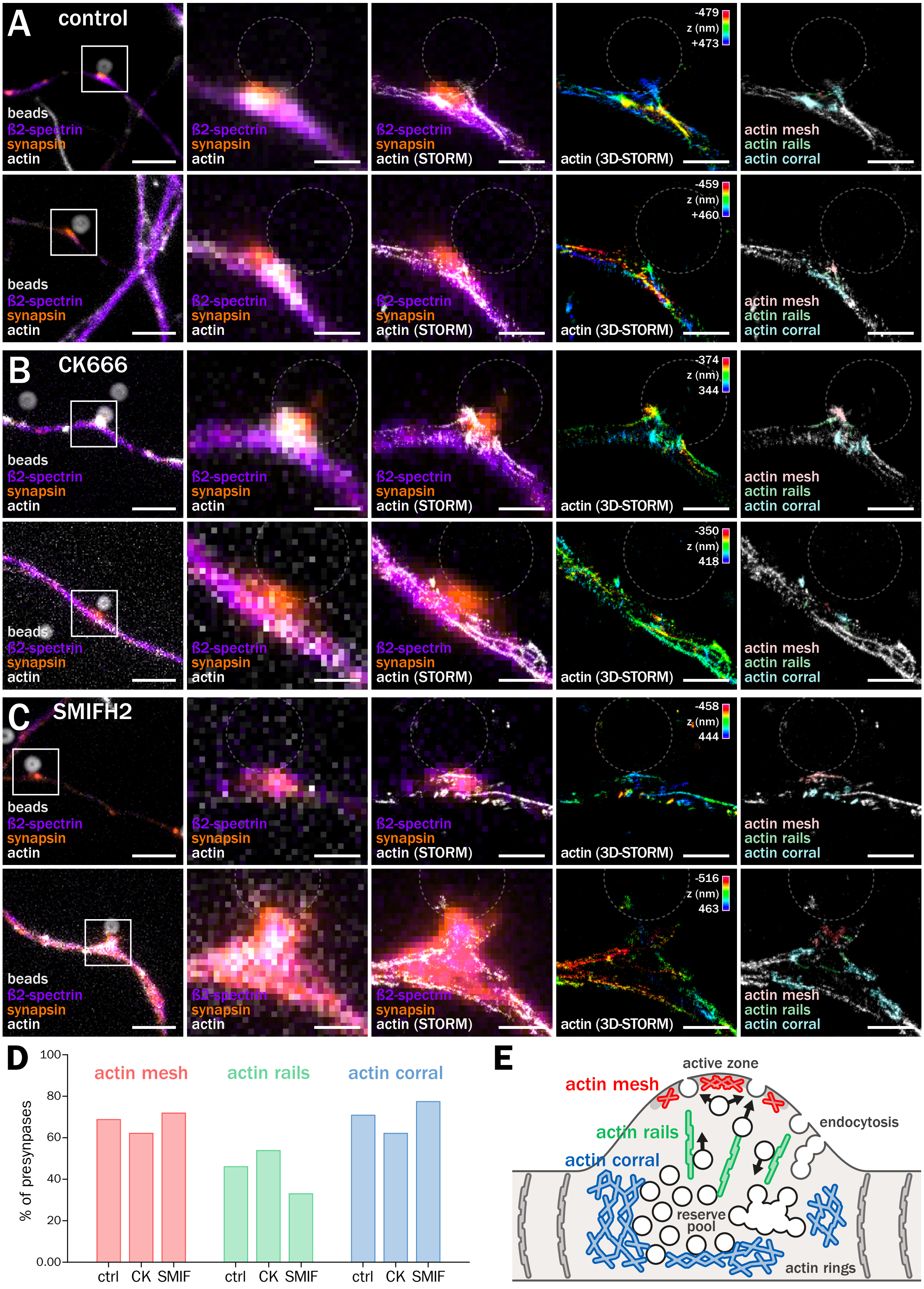
A
control
beads
ß2-spectrin
synapsin
actin
actin (STORM)
actin (3D-STORM)
z (nm)
-479
+473
-459
+460
actin mesh
actin rails
actin corral
B
CK666
-374
344
-350
418
C
SMIFH2
-458
444
-516
463
D
% of presynapses
100
80
60
40
20
0.00
ctrl
CK
SMIF
E
active zone
endocytosis
reserve pool
actin rings

#### Figure S1

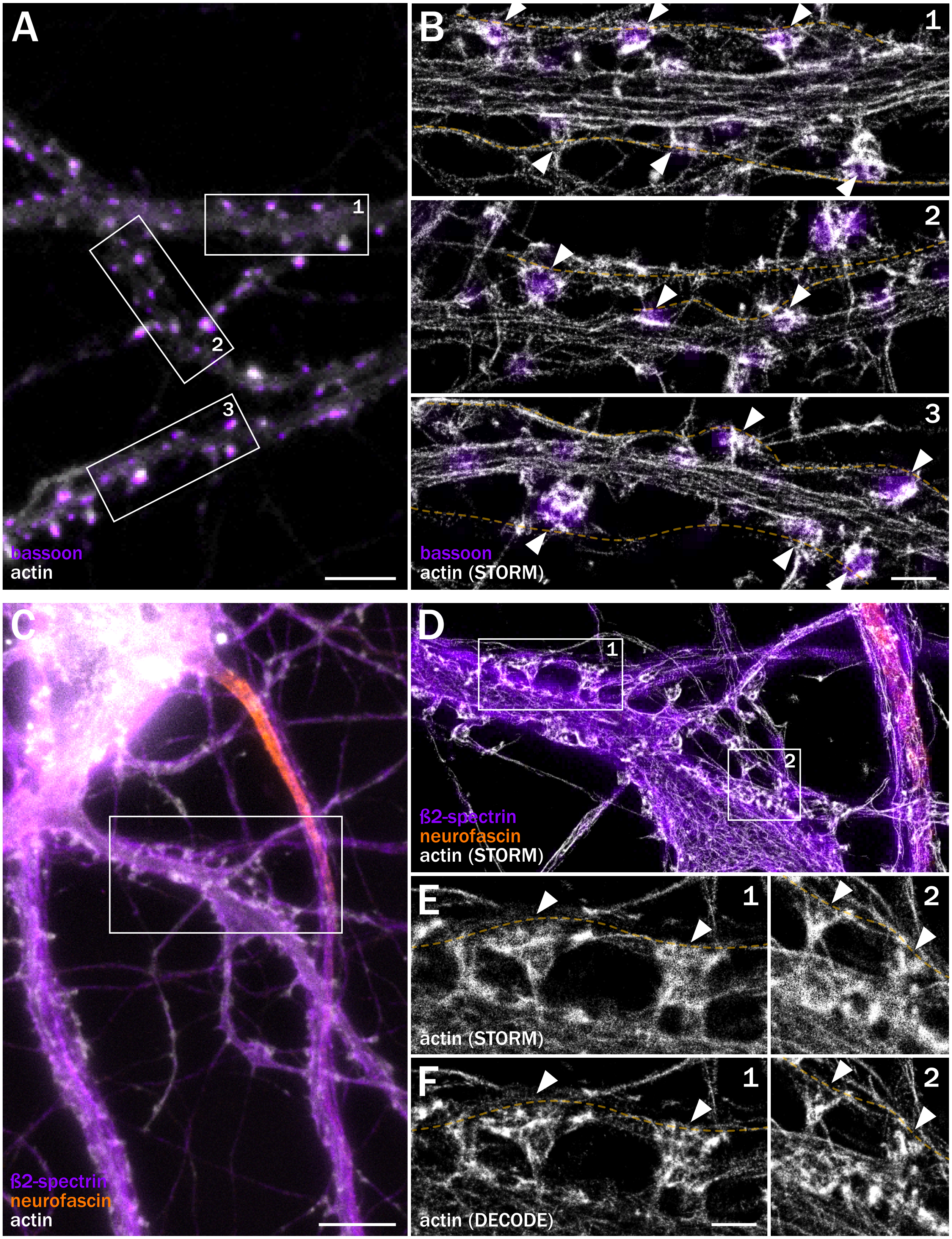
A
1
2
3
bassoon
actin
B
1
2
3
bassoon
actin (STORM)
C
ß2-spectrin
neurofascin
actin
D
1
2
ß2-spectrin
neurofascin
actin (STORM)
E
1
2
actin (STORM)
F
1
2
actin (DECODE)

### Figure S2

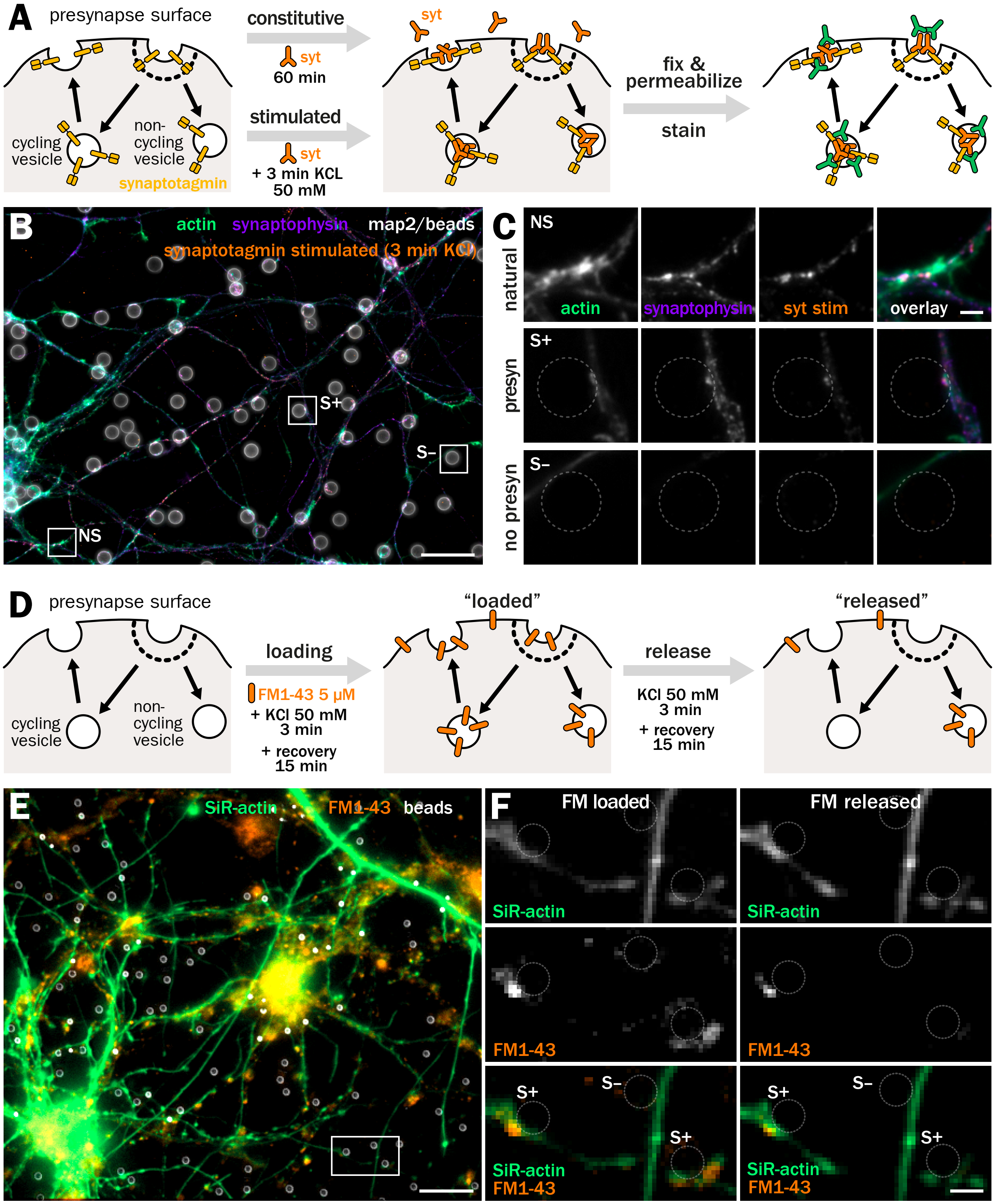
A
presynapse surface
cycling vesicle
non-cycling vesicle
synaptotagmin
constitutive
syt
60 min
stimulated
syt
+ 3 min KCL 50 mM
syt
fix & permeabilize
stain
B
actin
synaptophysin
map2/beads
synaptotagmin stimulated (3 min KCl)
S+
S–
NS
C
NS
natural
actin
synaptophysin
syt stim
overlay
S+
presyn
S–
no presyn
D
presynapse surface
cycling vesicle
non-cycling vesicle
loading
FM1-43 5 µM
+ KCl 50 mM 3 min
+ recovery 15 min
"loaded"
release
KCl 50 mM 3 min
+ recovery 15 min
"released"
E
SiR-actin
FM1-43
beads
F
FM loaded
FM released
SiR-actin
SiR-actin
FM1-43
FM1-43
S+
S–
S+
SiR-actin
FM1-43
S+
S–
S+
SiR-actin
FM1-43

#### Figure S3

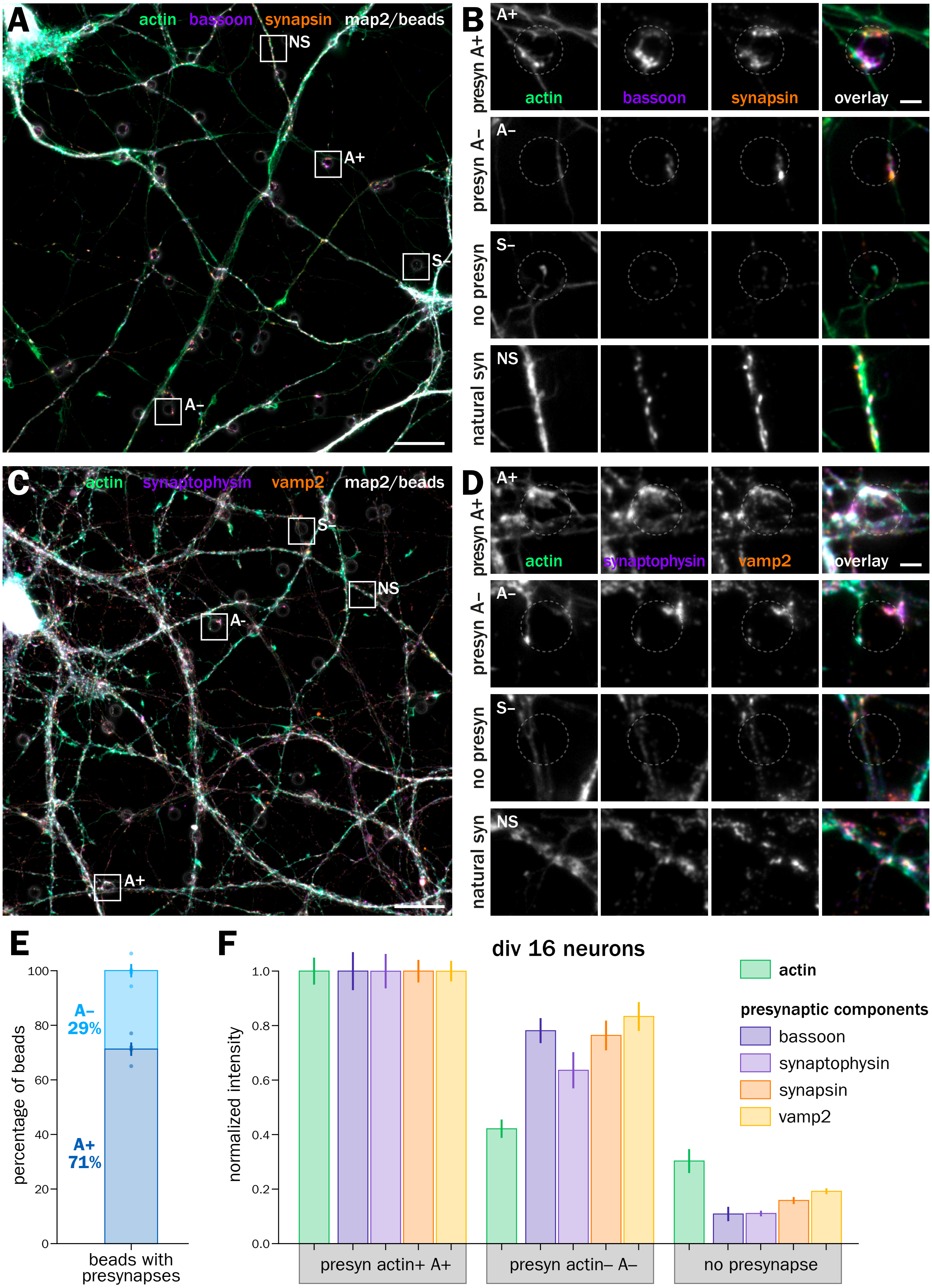
A
actin bassoon synapsin map2/beads
NS
A+
S–
A–
B
presyn A+
A+
actin
bassoon
synapsin
overlay
presyn A–
A–
no presyn
S–
natural syn
NS
C
actin synaptophysin vamp2 map2/beads
S–
NS
A–
A+
D
presyn A+
A+
actin
synaptophysin
vamp2
overlay
presyn A–
A–
no presyn
S–
natural syn
NS
E
percentage of beads
100
80
60
40
20
0
A–
29%
A+
71%
beads with presynapses
F
div 16 neurons
normalized intensity
1.0
0.8
0.6
0.4
0.2
0.0
presyn actin+ A+
presyn actin– A–
no presynapse
actin
presynaptic components
bassoon
synaptophysin
synapsin
vamp2

### Figure S4

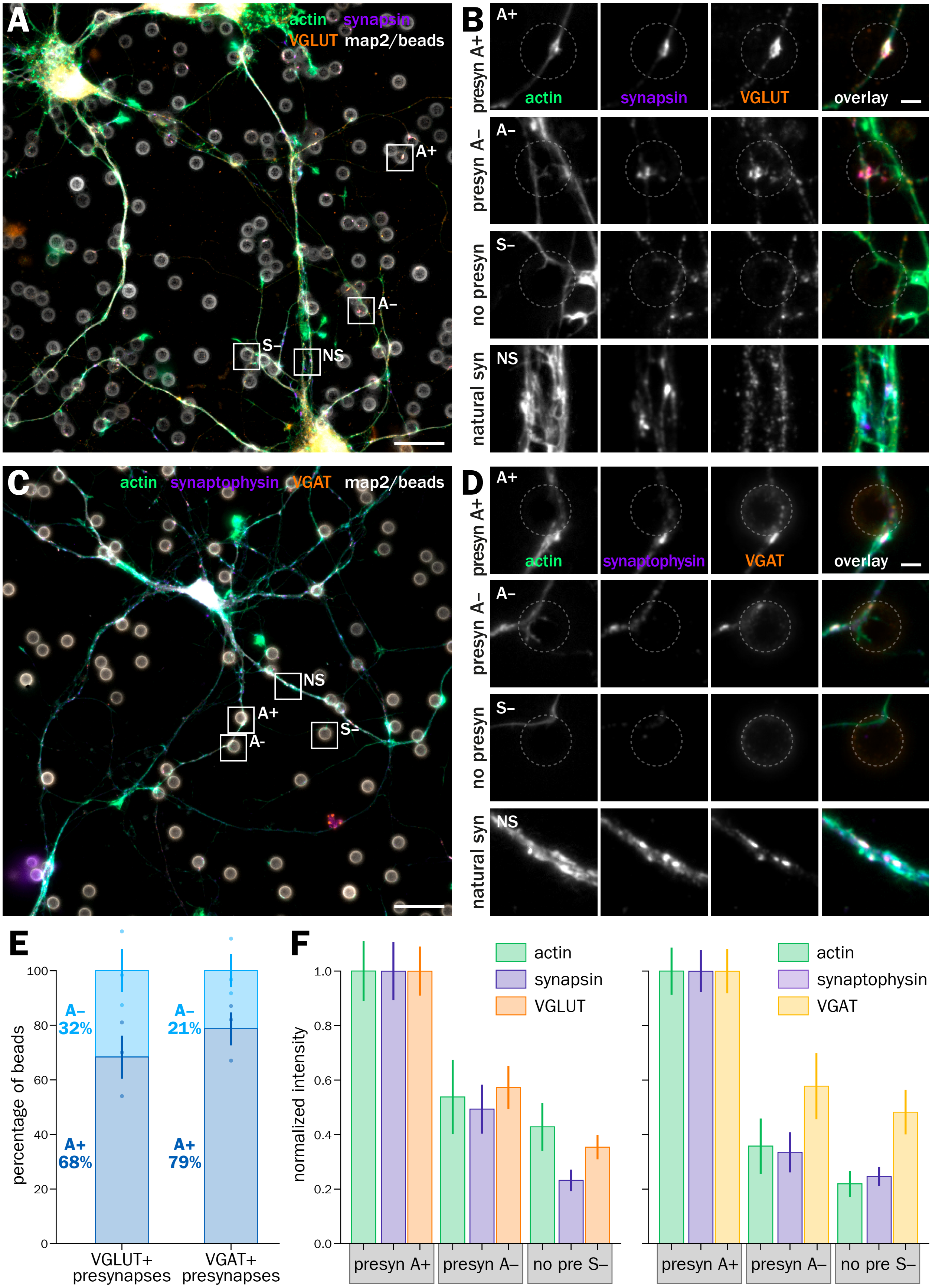
A
actin synapsin
VGLUT map2/beads
A+
A–
S–
NS
B
presyn A+
presyn A–
no presyn
natural syn
actin
synapsin
VGLUT
overlay
C
actin synaptophysin VGAT map2/beads
D
synaptophysin
VGAT
E
percentage of beads
A–
32%
A+
68%
21%
79%
VGLUT+
presynapses
VGAT+
presynapses
F
normalized intensity
presyn A+
presyn A–
no pre S–

#### Figure S5

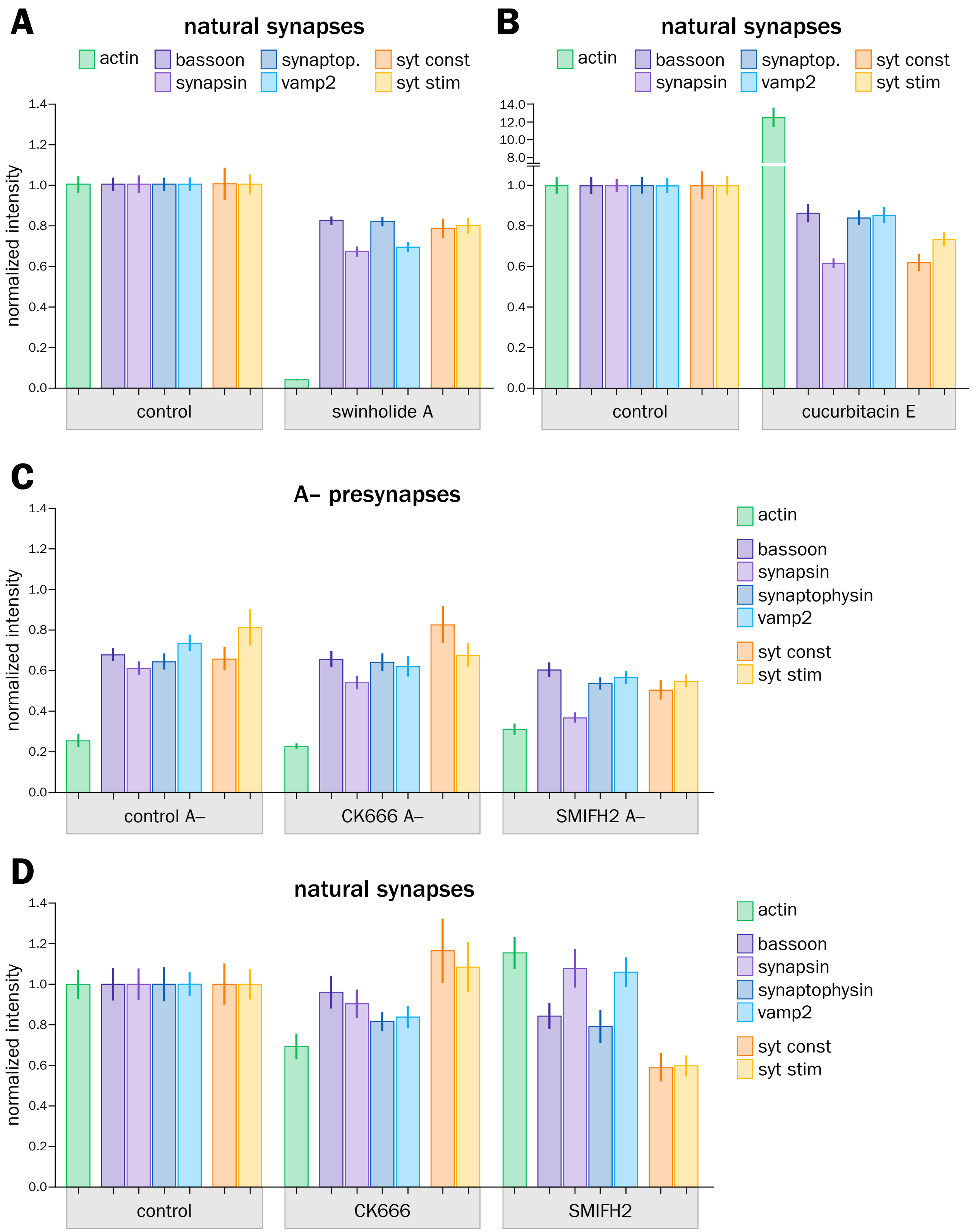
A
natural synapses
actin
bassoon
synapsin
synaptop.
vamp2
syt const
syt stim
normalized intensity
1.4
1.2
1.0
0.8
0.6
0.4
0.2
0.0
control
swinholide A
B
natural synapses
14.0
12.0
10.0
8.0
cucurbitacin E
C
A– presynapses
synaptophysin
control A–
CK666 A–
SMIFH2 A–
D
natural synapses
control
CK666
SMIFH2
