## Supplementary File 1 for "Distinct nano-structures support a multifunctional role of actin at presynapses"

Statistics for the Manuscript:
### Distinct nano-structures support a multifunctional role of actin at presynapses
Bingham et al., bioRxiv 2022

#### Statistics for Figure 1

| **Fig. 1G** | **Counts: bead-induced presynapses S+/S-** | | | | | |
|---|---|---|---|---|---|---|
| ***Condit 1*** | ***Condit 2*** | ***n points*** | ***N exp*** | ***Mean*** | ***SD*** | ***SEM*** |
| beads | S+ | 661 | 8 | 79,96 | 6,57 | 2,32 |
| beads | S- | 148 | 8 | 20,04 | 6,56 | 2,32 |

| **Fig. 1H** | **Intensities: Natural synapses vs bead-induced presynapses** | | | | | | | |
|---|---|---|---|---|---|---|---|---|
| ***Condit 1*** | ***Condit 2*** | ***n points*** | ***N exp*** | ***Mean*** | ***SD*** | ***SEM*** | ***Sig 1*** | ***against*** |
| bass | NS | 94 | 3 | 2,10 | 0,82 | 0,08 | *** | bass S+ |
| | S+ | 143 | 3 | 1,00 | 0,55 | 0,05 | | |
| | S- | 36 | 3 | 0,12 | 0,09 | 0,02 | *** | bass S+ |
| syn | NS | 191 | 4 | 1,92 | 1,28 | 0,09 | *** | syn S+ |
| | S+ | 364 | 4 | 1,00 | 0,62 | 0,03 | | |
| | S- | 70 | 4 | 0,24 | 0,21 | 0,02 | *** | syn S+ |
| syp | NS | 155 | 3 | 1,81 | 0,87 | 0,07 | *** | syp S+ |
| | S+ | 303 | 3 | 1,00 | 0,59 | 0,03 | | |
| | S- | 60 | 3 | 0,21 | 0,17 | 0,02 | *** | syp S+ |
| vamp | NS | 103 | 3 | 1,94 | 0,66 | 0,07 | *** | vamp S+ |
| | S+ | 226 | 3 | 1,00 | 0,57 | 0,04 | | |
| | S- | 35 | 3 | 0,20 | 0,16 | 0,03 | *** | vamp S+ |

| **Fig. 1I** | **Intensities: Natural synapses vs bead-induced presynapses** | | | | | | | |
|---|---|---|---|---|---|---|---|---|
| ***Condit 1*** | ***Condit 2*** | ***n points*** | ***N exp*** | ***Mean*** | ***SD*** | ***SEM*** | ***Sig 1*** | ***against*** |
| syt feed | NS | 128 | 3 | 1,61 | 1,81 | 0,16 | *** | syt feed S+ |
| | S+ | 295 | 3 | 1,00 | 1,00 | 0,06 | | |
| | S- | 294 | 3 | 0,22 | 0,39 | 0,02 | *** | syt feed S+ |
| syt KCl | NS | 151 | 3 | 1,36 | 1,03 | 0,08 | *** | syt KCl S+ |
| | S+ | 343 | 3 | 1,00 | 0,85 | 0,05 | | |
| | S- | 340 | 3 | 0,32 | 0,25 | 0,01 | *** | syt KCl S+ |

### Statistics for Figure 2

| Fig. 2E | Counts: bead-induced presynapses A+/A- | | | | | |
|---|---|---|---|---|---|---|
| *Condit 1* | *Condit 2* | *n points* | *N exp* | *Mean* | *SD* | *SEM* |
| all beads | A+ | 437 | 7 | 53,08 | 6,03 | 2,13 |
| all beads | A- | 224 | 7 | 26,89 | 7,12 | 2,52 |
| all beads | S- | 148 | 7 | 20,05 | 6,56 | 2,32 |
| beads w/ | A+ | 437 | 7 | 66,59 | 7,95 | 2,81 |
| beads w/ | A- | 224 | 7 | 33,41 | 7,95 | 2,81 |

| Fig. 2F | Intensities: bead-induced presynapses A+/A- | | | | | | | | | |
|---|---|---|---|---|---|---|---|---|---|---|
| *Condit 1* | *Condit 2* | *n points* | *N exp* | *Mean* | *SD* | *SEM* | *Sig 1* | *against* | *Sig 2* | *against* |
| actin | A+ | 437 | 7 | 1,00 | 1,25 | 0,06 | | | | |
| | A- | 224 | 7 | 0,25 | 0,43 | 0,03 | *** | actin A+ | ns | actin S- |
| | S- | 148 | 7 | 0,29 | 0,34 | 0,03 | *** | actin A+ | | |
| bass | A+ | 210 | 6 | 1,00 | 0,52 | 0,04 | | | | |
| | A- | 104 | 6 | 0,74 | 0,31 | 0,03 | *** | bass A+ | *** | bass S- |
| | S- | 75 | 6 | 0,10 | 0,07 | 0,01 | *** | bass A+ | | |
| syn | A+ | 357 | 7 | 1,00 | 0,53 | 0,03 | | | | |
| | A- | 178 | 7 | 0,70 | 0,40 | 0,03 | *** | syn A+ | *** | syn S- |
| | S- | 108 | 7 | 0,20 | 0,18 | 0,02 | *** | syn A+ | | |
| syp | A+ | 294 | 6 | 1,00 | 0,54 | 0,03 | | | | |
| | A- | 140 | 6 | 0,66 | 0,37 | 0,03 | *** | syp A+ | *** | syp S- |
| | S- | 101 | 6 | 0,16 | 0,14 | 0,01 | *** | syp A+ | | |
| vamp | A+ | 232 | 6 | 1,00 | 0,49 | 0,03 | | | | |
| | A- | 124 | 6 | 0,65 | 0,35 | 0,03 | *** | vamp A+ | *** | vamp S- |
| | S- | 76 | 6 | 0,18 | 0,13 | 0,01 | *** | vamp A+ | | |

### Statistics for Figure 3

| Fig. 3E | Syt endocytosis: bead-induced presynapses A+/A- | | | | | | | | | |
|---|---|---|---|---|---|---|---|---|---|---|
| ***Condit 1*** | ***Condit 2*** | ***n points*** | ***N exp*** | ***Mean*** | ***SD*** | ***SEM*** | ***Sig 1*** | ***against*** | ***Sig 2*** | ***against*** |
| syt feed | A+ | 145 | 3 | 1,00 | 0,82 | 0,07 | | | | |
| | A- | 81 | 3 | 0,82 | 0,72 | 0,08 | ns | syt feed A+ | *** | syt feed S- |
| | S- | 112 | 3 | 0,30 | 0,29 | 0,03 | *** | syt feed A+ | | |
| syt KCl | A+ | 195 | 3 | 1,00 | 0,55 | 0,04 | | | | |
| | A- | 62 | 3 | 0,72 | 0,26 | 0,03 | *** | syt KCl A+ | *** | syt KCl S- |
| | S- | 131 | 3 | 0,28 | 0,18 | 0,02 | *** | syt KCl A+ | | |

| Fig. 3G | FM cycling: bead-induced presynapses A+/A- | | | | | | | | | | | |
|---|---|---|---|---|---|---|---|---|---|---|---|---|
| ***Condit 1*** | ***Condit 2*** | ***n points*** | ***N exp*** | ***Mean*** | ***SD*** | ***SEM*** | ***Sig 1*** | ***against*** | ***Sig 2*** | ***against*** | ***Sig 3*** | ***against*** |
| loaded | A+ | 60 | 3 | 1,00 | 0,58 | 0,07 | | | | | | |
| | A- | 28 | 3 | 0,71 | 0,35 | 0,07 | ** | load A+ | *** | load S- | | |
| | S- | 89 | 3 | 0,27 | 0,25 | 0,03 | *** | load A+ | | | | |
| releasec | A+ | 38 | 2 | 0,38 | 0,19 | 0,03 | | | | | *** | load A+ |
| | A- | 11 | 2 | 0,32 | 0,14 | 0,04 | ns | rel A+ | ns | rel S- | ** | load A- |
| | S- | 48 | 2 | 0,12 | 0,08 | 0,01 | ** | rel A+ | | | ns | load S- |

### Statistics for Figure 4

| **Fig. 4D** | **Swinholide A effects** | | | | | | | | | |
|---|---|---|---|---|---|---|---|---|---|---|
| ***Condit 1*** | ***Condit 2*** | ***n points*** | ***N exp*** | ***Mean*** | ***SD*** | ***SEM*** | ***Sig 1*** | ***against*** | ***Sig 2*** | ***against*** |
| actin | A+ | 402 | 8 | 1,00 | 1,27 | 0,06 | | | | |
| | A- | 193 | 8 | 0,19 | 0,52 | 0,04 | | | | |
| | swin A- | 500 | 8 | 0,02 | 0,18 | 0,01 | *** | actin A+ | ** | actin A- |
| bass | A+ | 105 | 3 | 1,00 | 0,55 | 0,05 | | | | |
| | A- | 38 | 3 | 0,71 | 0,26 | 0,04 | | | | |
| | swin A- | 101 | 3 | 0,69 | 0,31 | 0,03 | *** | bass A+ | ns | bass A- |
| syn | A+ | 252 | 4 | 1,00 | 0,57 | 0,04 | | | | |
| | A- | 112 | 4 | 0,67 | 0,42 | 0,04 | | | | |
| | swin A- | 224 | 4 | 0,67 | 0,41 | 0,03 | *** | syn A+ | ns | syn A- |
| syp | A+ | 256 | 3 | 1,00 | 0,55 | 0,03 | | | | |
| | A- | 124 | 3 | 0,66 | 0,38 | 0,03 | | | | |
| | swin A- | 312 | 3 | 0,73 | 0,48 | 0,03 | *** | syp A+ | ns | syp A- |
| vamp | A+ | 150 | 3 | 1,00 | 0,51 | 0,04 | | | | |
| | A- | 76 | 3 | 0,58 | 0,31 | 0,04 | | | | |
| | swin A- | 125 | 3 | 0,61 | 0,28 | 0,03 | *** | vamp A+ | ns | vamp A- |
| syt feed | A+ | 145 | 3 | 1,00 | 0,82 | 0,07 | | | | |
| | A- | 79 | 3 | 0,76 | 0,59 | 0,07 | | | | |
| | swin A- | 268 | 3 | 0,79 | 0,73 | 0,04 | ** | syt feed A+ | ns | syt feed A- |
| syt KCl | A+ | 195 | 3 | 1,00 | 0,55 | 0,04 | | | | |
| | A- | 62 | 3 | 0,72 | 0,26 | 0,03 | | | | |
| | swin A- | 206 | 3 | 0,71 | 0,38 | 0,03 | *** | syt KCl A+ | ns | syt KCl A- |

| **Fig. 4E** | **Cucurbitacin E effects** | | | | | | | | | |
|---|---|---|---|---|---|---|---|---|---|---|
| ***Condit 1*** | ***Condit 2*** | ***n points*** | ***N exp*** | ***Mean*** | ***SD*** | ***SEM*** | ***Sig 1*** | ***against*** | ***Sig 2*** | ***against*** |
| actin | A+ | 475 | 8 | 1,00 | 0,84 | 0,04 | | | | |
| | A- | 174 | 8 | 0,28 | 0,43 | 0,03 | | | | |
| | cuc A+ | 409 | 8 | 7,30 | 8,77 | 0,43 | *** | actin A+ | *** | actin A- |
| bass | A+ | 105 | 2 | 1,00 | 0,45 | 0,04 | | | | |
| | A- | 36 | 2 | 0,75 | 0,30 | 0,05 | | | | |
| | cuc A+ | 96 | 2 | 0,83 | 0,43 | 0,04 | *** | bass A+ | ns | bass A- |
| syn | A+ | 203 | 4 | 1,00 | 0,50 | 0,04 | | | | |
| | A- | 62 | 4 | 0,67 | 0,29 | 0,04 | | | | |
| | cuc A+ | 229 | 4 | 0,68 | 0,35 | 0,02 | *** | syn A+ | ns | syn A- |
| syp | A+ | 334 | 7 | 1,00 | 0,73 | 0,04 | | | | |
| | A- | 125 | 7 | 0,71 | 0,59 | 0,05 | | | | |
| | cuc A+ | 310 | 7 | 0,86 | 0,69 | 0,04 | ** | syp A+ | * | syp A- |
| vamp | A+ | 189 | 4 | 1,00 | 0,59 | 0,04 | | | | |
| | A- | 45 | 4 | 0,63 | 0,36 | 0,05 | | | | |
| | cuc A+ | 167 | 4 | 0,89 | 0,40 | 0,03 | * | vamp A+ | *** | vamp A- |
| syt feed | A+ | 145 | 3 | 1,00 | 0,82 | 0,07 | | | | |
| | A- | 79 | 3 | 0,76 | 0,59 | 0,07 | | | | |
| | cuc A+ | 141 | 3 | 0,77 | 0,93 | 0,08 | * | syt feed A+ | ns | syt feed A- |
| syt KCl | A+ | 195 | 3 | 1,00 | 0,55 | 0,04 | | | | |
| | A- | 62 | 3 | 0,72 | 0,26 | 0,03 | | | | |
| | cuc A+ | 138 | 3 | 0,69 | 0,43 | 0,04 | *** | syt KCl A+ | ns | syt KCl A- |

### Statistics for Figure 5

| Fig. 5D | CK 666 and SMIFH2 effects on A+ presynapses | | | | | | | |
|---|---|---|---|---|---|---|---|---|
| ***Condit 1*** | ***Condit 2*** | ***n points*** | ***N exp*** | ***Mean*** | ***SD*** | ***SEM*** | ***Sig 1*** | ***against*** |
| actin | A+ | 887 | 8 | 1,00 | 1,07 | 0,04 | | |
| | CK 666 A+ | 817 | 8 | 0,95 | 1,19 | 0,04 | ns | actin A+ |
| | SMIFH2 A+ | 695 | 8 | 1,35 | 1,55 | 0,06 | *** | actin A+ |
| bass | A+ | 356 | 4 | 1,00 | 0,67 | 0,04 | | |
| | CK 666 A+ | 344 | 4 | 0,89 | 0,67 | 0,04 | ** | bass A+ |
| | SMIFH2 A+ | 259 | 4 | 0,86 | 0,66 | 0,04 | *** | bass A+ |
| syn | A+ | 356 | 4 | 1,00 | 0,69 | 0,04 | | |
| | CK 666 A+ | 344 | 4 | 0,77 | 0,71 | 0,04 | *** | syn A+ |
| | SMIFH2 A+ | 259 | 4 | 0,80 | 0,86 | 0,05 | *** | syn A+ |
| syp | A+ | 315 | 4 | 1,00 | 0,61 | 0,03 | | |
| | CK 666 A+ | 275 | 4 | 1,04 | 0,84 | 0,05 | ns | syp A+ |
| | SMIFH2 A+ | 260 | 4 | 0,71 | 0,55 | 0,03 | *** | syp A+ |
| vamp | A+ | 324 | 4 | 1,00 | 0,57 | 0,03 | | |
| | CK 666 A+ | 331 | 4 | 1,07 | 0,67 | 0,04 | ns | vamp A+ |
| | SMIFH2 A+ | 287 | 4 | 0,89 | 0,65 | 0,04 | * | vamp A+ |
| syt feed | A+ | 211 | 3 | 1,00 | 0,98 | 0,07 | | |
| | CK 666 A+ | 141 | 3 | 1,05 | 1,08 | 0,09 | ns | syt feed A+ |
| | SMIFH2 A+ | 194 | 3 | 0,55 | 0,40 | 0,03 | *** | syt feed A+ |
| syt KCl | A+ | 228 | 3 | 1,00 | 0,70 | 0,05 | | |
| | CK A+ | 187 | 3 | 1,08 | 0,94 | 0,07 | ns | syt KCl A+ |
| | SMIFH2 A+ | 168 | 3 | 0,72 | 0,46 | 0,04 | *** | syt KCl A+ |

### Statistics for Figure 7

| Fig. 7D | Actin nano-structures after CK666 and SMIFH2 treatments | | | | | | | |
|---|---|---|---|---|---|---|---|---|
| ***Condit 1*** | ***Condit 2*** | ***Found*** | ***Not Found*** | ***Total F+NF*** | ***%*** | ***ND*** | ***Total*** | ***N exp*** |
| mesh | control | 47 | 19 | 66 | 71,21 | 2 | 68 | 5 |
| | CK 666 | 20 | 12 | 32 | 62,50 | 0 | 32 | 3 |
| | SMIFH2 | 14 | 4 | 18 | 77,78 | 0 | 18 | 2 |
| rails | control | 26 | 30 | 56 | 46,43 | 12 | 68 | 5 |
| | CK 666 | 13 | 11 | 24 | 54,17 | 8 | 32 | 3 |
| | SMIFH2 | 5 | 10 | 15 | 33,33 | 3 | 18 | 2 |
| corral | control | 47 | 21 | 68 | 69,12 | 0 | 68 | 5 |
| | CK 666 | 20 | 12 | 32 | 62,50 | 0 | 32 | 3 |
| | SMIFH2 | 13 | 5 | 18 | 72,22 | 0 | 18 | 2 |

### Statistics for Figure S3

| Fig. S3E | Div 14 counts: induced presynapses A+/A- | | | | | |
|---|---|---|---|---|---|---|
| ***Condit 1*** | ***Condit 2*** | ***n points*** | ***N exp*** | ***Mean*** | ***SD*** | ***SEM*** |
| beads | A+ | 129 | 3 | 71,25 | 4,92 | 2,46 |
| beads | A- | 54 | 3 | 28,75 | 4,92 | 2,46 |

| Fig. S3F | Div 14 intensities: bead-induced presynapses A+/A- | | | | | | | | | |
|---|---|---|---|---|---|---|---|---|---|---|
| ***Condit 1*** | ***Condit 2*** | ***n points*** | ***N exp*** | ***Mean*** | ***SD*** | ***SEM*** | ***Sig 1*** | ***against*** | ***Sig 2*** | ***against*** |
| actin | A+ | 130 | 3 | 1,00 | 0,57 | 0,05 | | | | |
| | A- | 68 | 3 | 0,42 | 0,28 | 0,03 | *** | actin A+ | ns | actin S- |
| | S- | 60 | 3 | 0,30 | 0,34 | 0,04 | *** | actin A+ | | |
| bass | A+ | 51 | 2 | 1,00 | 0,50 | 0,07 | | | | |
| | A- | 34 | 2 | 0,78 | 0,27 | 0,05 | * | bass A+ | *** | bass S- |
| | S- | 11 | 2 | 0,11 | 0,09 | 0,09 | *** | bass A+ | | |
| syn | A+ | 51 | 2 | 1,00 | 0,45 | 0,06 | | | | |
| | A- | 18 | 2 | 0,64 | 0,28 | 0,07 | *** | syn A+ | *** | syn S- |
| | S- | 26 | 2 | 0,11 | 0,05 | 0,01 | *** | syn A+ | | |
| syp | A+ | 79 | 2 | 1,00 | 0,37 | 0,04 | | | | |
| | A- | 36 | 2 | 0,76 | 0,33 | 0,05 | *** | syp A+ | *** | syp S- |
| | S- | 34 | 2 | 0,16 | 0,07 | 0,01 | *** | syp A+ | | |
| vamp | A+ | 78 | 2 | 1,00 | 0,33 | 0,04 | | | | |
| | A- | 36 | 2 | 0,83 | 0,32 | 0,05 | * | vamp A+ | *** | vamp S- |
| | S- | 34 | 2 | 0,19 | 0,06 | 0,01 | *** | vamp A+ | | |

### Statistics for Figure S4

| **Fig. S4E** | **GLUT/GABA counts: presynapses A+/A-** | | | | | |
|---|---|---|---|---|---|---|
| **Condit 1** | **Condit 2** | **n points** | **N exp** | **Mean** | **SD** | **SEM** |
| VGLUT | A+ | 185 | 3 | 68,33 | 13,58 | 7,84 |
| | A- | 75 | 3 | 31,67 | 13,58 | 7,84 |
| VGAT | A+ | 101 | 3 | 78,67 | 10,41 | 6,01 |
| | A- | 33 | 3 | 21,33 | 10,41 | 6,01 |

| **Fig. S4F** | **GLUT/GABA intensities: presynapses A+/A-** | | | | | | | | | |
|---|---|---|---|---|---|---|---|---|---|---|
| **Condit 1** | **Condit 2** | **n points** | **N exp** | **Mean** | **SD** | **SEM** | **Sig 1** | **against** | **Sig 2** | **against** |
| VGLUT | actin A+ | 185 | 3 | 1,00 | 1,50 | 0,11 | | | | |
| | actin A- | 75 | 3 | 0,54 | 1,18 | 0,14 | * | actin A+ | ns | actin S- |
| | actin S- | 40 | 3 | 0,43 | 0,56 | 0,09 | * | actin A+ | | |
| VGLUT | syn A+ | 185 | 3 | 1,00 | 1,45 | 0,11 | | | | |
| | syn A- | 75 | 3 | 0,49 | 0,77 | 0,09 | ** | syn A+ | ns | syn S- |
| | syn S- | 40 | 3 | 0,23 | 0,25 | 0,04 | *** | syn A+ | | |
| VGLUT | VGLUT A+ | 185 | 3 | 1,00 | 1,23 | 0,09 | | | | |
| | VGLUT A- | 75 | 3 | 0,57 | 0,69 | 0,08 | ** | VGLUT A+ | ns | VGLUT S- |
| | VGLUT S- | 40 | 3 | 0,35 | 0,28 | 0,04 | ** | VGLUT A+ | | |
| VGAT | actin A+ | 101 | 3 | 1,00 | 0,87 | 0,09 | | | | |
| | actin A- | 33 | 3 | 0,36 | 0,56 | 0,10 | ** | actin A+ | ns | actin S- |
| | actin S- | 20 | 3 | 0,22 | 0,22 | 0,05 | ** | actin A+ | | |
| VGAT | syp A+ | 101 | 3 | 1,00 | 0,77 | 0,08 | | | | |
| | syp A- | 33 | 3 | 0,34 | 0,42 | 0,07 | *** | syp A+ | ns | syp S- |
| | syp S- | 20 | 3 | 0,25 | 0,16 | 0,03 | *** | syp A+ | | |
| VGAT | VGAT A+ | 101 | 3 | 1,00 | 0,82 | 0,08 | | | | |
| | VGAT A- | 33 | 3 | 0,58 | 0,70 | 0,12 | * | VGAT A+ | ns | VGAT S- |
| | VGAT S- | 20 | 3 | 0,48 | 0,37 | 0,08 | * | VGAT A+ | | |

### Statistics for Figure S5A-B

| Fig. S5A | Swinholide A effects on natural synapses | | | | | | | |
|---|---|---|---|---|---|---|---|---|
| ***Condit 1*** | ***Condit 2*** | ***n points*** | ***N exp*** | ***Mean*** | ***SD*** | ***SEM*** | ***Sig 1*** | ***against*** |
| actin | control | 675 | 10 | 1,00 | 1,07 | 0,04 | | |
| | swin | 673 | 10 | 0,04 | 0,08 | 0,00 | *** | control |
| bass | control | 94 | 3 | 1,00 | 0,32 | 0,03 | | |
| | swin | 125 | 3 | 0,82 | 0,23 | 0,02 | *** | control |
| syn | control | 191 | 4 | 1,00 | 0,59 | 0,04 | | |
| | swin | 220 | 4 | 0,67 | 0,38 | 0,03 | *** | control |
| syp | control | 266 | 3 | 1,00 | 0,53 | 0,03 | | |
| | swin | 385 | 3 | 0,82 | 0,47 | 0,02 | *** | control |
| vamp | control | 101 | 3 | 1,00 | 0,34 | 0,03 | | |
| | swin | 107 | 3 | 0,69 | 0,25 | 0,02 | *** | control |
| syt feed | control | 213 | 3 | 1,00 | 1,15 | 0,08 | | |
| | swin | 207 | 3 | 0,78 | 0,69 | 0,05 | * | control |
| syt KCl | control | 223 | 3 | 1,00 | 0,72 | 0,05 | | |
| | swin | 232 | 3 | 0,80 | 0,59 | 0,04 | * | control |

| Fig. S5B | Cucurbitacin E effects on natural synapses | | | | | | | |
|---|---|---|---|---|---|---|---|---|
| ***Condit 1*** | ***Condit 2*** | ***n points*** | ***N exp*** | ***Mean*** | ***SD*** | ***SEM*** | ***Sig 1*** | ***against*** |
| actin | control | 490 | 10 | 1,00 | 1,06 | 0,04 | | |
| | cuc | 662 | 10 | 12,52 | 24,46 | 1,11 | *** | control |
| bass | control | 79 | 3 | 1,00 | 0,38 | 0,04 | | |
| | cuc | 81 | 3 | 0,86 | 0,40 | 0,04 | * | control |
| syn | control | 113 | 3 | 1,00 | 0,34 | 0,03 | | |
| | cuc | 121 | 3 | 0,62 | 0,26 | 0,02 | *** | control |
| syp | control | 71 | 3 | 1,00 | 0,34 | 0,04 | | |
| | cuc | 78 | 3 | 0,84 | 0,31 | 0,04 | *** | control |
| vamp | control | 71 | 3 | 1,00 | 0,32 | 0,04 | | |
| | cuc | 78 | 3 | 0,85 | 0,36 | 0,04 | *** | control |
| syt feed | control | 223 | 3 | 1,00 | 1,10 | 0,07 | | |
| | cuc | 217 | 3 | 0,62 | 0,62 | 0,04 | *** | control |
| syt KCl | control | 255 | 3 | 1,00 | 0,72 | 0,05 | | |
| | cuc | 185 | 3 | 0,74 | 0,47 | 0,03 | ** | control |

### Statistic for Figure S5C-D

| Fig. S5C | CK666 and SMIFH2 effects on A- presynapses | | | | | | | |
|---|---|---|---|---|---|---|---|---|
| ***Condit 1*** | ***Condit 2*** | ***n points*** | ***N exp*** | ***Mean*** | ***SD*** | ***SEM*** | ***Sig 1*** | ***against*** |
| actin | A- | 326 | 8 | 0,26 | 0,58 | 0,03 | | |
| | CK 666 A | 293 | 8 | 0,23 | 0,25 | 0,01 | ns | actin A- |
| | SMIFH2 A- | 283 | 8 | 0,31 | 0,47 | 0,03 | ns | actin A- |
| bass | A- | 146 | 4 | 0,68 | 0,38 | 0,03 | | |
| | CK 666 A | 135 | 4 | 0,66 | 0,46 | 0,04 | ns | bass A- |
| | SMIFH2 A- | 109 | 4 | 0,61 | 0,37 | 0,04 | ns | bass A- |
| syn | A- | 147 | 4 | 0,61 | 0,40 | 0,03 | | |
| | CK 666 A | 134 | 4 | 0,54 | 0,39 | 0,03 | ns | syn A- |
| | SMIFH2 A- | 109 | 4 | 0,37 | 0,27 | 0,03 | *** | syn A- |
| syp | A- | 99 | 4 | 0,74 | 0,41 | 0,04 | | |
| | CK 666 A | 140 | 4 | 0,64 | 0,52 | 0,04 | ns | syp A- |
| | SMIFH2 A- | 100 | 4 | 0,54 | 0,30 | 0,03 | ** | syp A- |
| vamp | A- | 108 | 4 | 0,65 | 0,41 | 0,04 | | |
| | CK 666 A- | 110 | 4 | 0,62 | 0,52 | 0,05 | ns | vamp A- |
| | SMIFH2 A- | 113 | 4 | 0,57 | 0,34 | 0,03 | ns | vamp A- |
| syt feed | A- | 84 | 3 | 0,66 | 0,53 | 0,06 | | |
| | CK 666 A | 69 | 3 | 0,83 | 0,76 | 0,09 | ns | syt feed A- |
| | SMIFH2 A- | 111 | 3 | 0,51 | 0,50 | 0,05 | ns | syt feed A- |
| syt KCl | A- | 114 | 3 | 0,81 | 0,96 | 0,09 | | |
| | CK 666 A- | 79 | 3 | 0,68 | 0,53 | 0,06 | ns | syt KCl A- |
| | SMIFH2 A- | 107 | 3 | 0,55 | 0,33 | 0,03 | *** | syt KCl A- |

| Fig. S5D | CK666 and SMIFH2 effects on natural synapses | | | | | | | |
|---|---|---|---|---|---|---|---|---|
| ***Condit 1*** | ***Condit 2*** | ***n points*** | ***N exp*** | ***Mean*** | ***SD*** | ***SEM*** | ***Sig 1*** | ***against*** |
| actin | control | 179 | 2 | 1,00 | 0,97 | 0,07 | | |
| | CK666 | 182 | 2 | 0,69 | 0,83 | 0,06 | ** | control |
| bass | control | 137 | 3 | 1,00 | 0,94 | 0,08 | | |
| | CK666 | 144 | 3 | 0,96 | 0,97 | 0,08 | ns | control |
| syn | control | 137 | 3 | 1,00 | 0,91 | 0,08 | | |
| | CK666 | 144 | 3 | 0,90 | 0,84 | 0,07 | ns | control |
| syp | control | 138 | 2 | 1,00 | 0,99 | 0,08 | | |
| | CK666 | 139 | 2 | 0,82 | 0,56 | 0,05 | ns | control |
| vamp | control | 143 | 3 | 1,00 | 0,72 | 0,06 | | |
| | CK666 | 149 | 3 | 0,84 | 0,68 | 0,06 | ns | control |
| syt feed | control | 128 | 3 | 1,00 | 1,16 | 0,10 | | |
| | CK666 | 121 | 3 | 1,17 | 1,74 | 0,16 | ns | control |
| syt KCl | control | 143 | 3 | 1,00 | 0,90 | 0,08 | | |
| | CK666 | 129 | 3 | 1,09 | 1,41 | 0,12 | ns | control |
| actin | control | 179 | 2 | 1,00 | 0,97 | 0,07 | | |
| | SMIFH2 | 184 | 2 | 1,16 | 1,07 | 0,08 | ns | control |
| bass | control | 137 | 3 | 1,00 | 0,94 | 0,08 | | |
| | SMIFH2 | 142 | 3 | 0,84 | 0,77 | 0,06 | ns | control |
| syn | control | 137 | 3 | 1,00 | 0,91 | 0,08 | | |
| | SMIFH2 | 145 | 3 | 1,08 | 1,15 | 0,10 | ns | control |
| syp | control | 138 | 2 | 1,00 | 0,99 | 0,08 | | |
| | SMIFH2 | 140 | 2 | 0,79 | 0,96 | 0,08 | ns | control |
| vamp | control | 143 | 3 | 1,00 | 0,72 | 0,06 | | |
| | SMIFH2 | 145 | 3 | 1,06 | 0,88 | 0,07 | ns | control |
| syt feed | control | | 3 | 1,00 | 1,16 | 0,10 | | |
| | SMIFH2 | 114 | 3 | 0,59 | 0,74 | 0,07 | ** | control |
| syt KCl | control | | 3 | 1,00 | 0,90 | 0,08 | | |
| | SMIFH2 | 112 | 3 | 0,60 | 0,54 | 0,05 | ** | control |
